# Learning of new associations invokes a major change in modulations of cortical beta oscillations in human adults

**DOI:** 10.1101/2022.05.02.490325

**Authors:** Anna Pavlova, Nikita Tyulenev, Vera Tretyakova, Valeriya Skavronskaya, Anastasia Nikolaeva, Andrey Prokofyev, Tatiana Stroganova, Boris Chernyshev

## Abstract

Large-scale cortical beta (β) oscillations have been implicated in the learning processes but their exact role is debated. We explored the dynamics of β-oscillations while 25 adult participants learned, through trial and error, novel associations between four auditory pseudowords and movements of four body extremities. We used MEG to evaluate learning-induced changes in beta modulation accompanying cue-triggered movements.

Our findings showed that spatial-temporal characteristics of movement-related β-oscillations underwent a major transition as learning proceeded. Early in learning, suppression of β-power in multiple cortical areas occurred long before movement initiation and sustained throughout the whole behavioral trial. As learning advanced and task performance reached asymptote, βsuppression was replaced by a widespread and prolonged rise in β-power. The β-power rise started shortly after the initiation of correct motor response and mainly comprised the prefrontal and medial temporal regions of the left hemisphere. This post-decision β-power predicted trial-by-trial response times (RT) at both stages of learning (before and after the rules become familiar) but in opposite ways. When a subject started to acquire associative rules and gradually improved task performance, a decrease in RT was correlated with the increase in the post-decision β-band power. Repeatedly correct implementation of the learned rules reversed this correlation in the opposite direction with faster (more confident) responses associated with the weaker post-decision β-band synchronization.

Our findings suggest that maximal beta activity is pertinent to a distinct stage of learning and may serve to strengthen the newly learned association in a distributed memory network.

## 1. Introduction

Numerous neuroimaging studies explored a relationship between processing of action language and activation of cortical areas related to motor functions. Converging findings revealed that action-related words trigger activation of higher-order cortical areas related to both perceptual processing and motor planning (premotor and supplementary motor regions) (Moseley & Pulvermüller, 2014; Pavlova et al., 2019; Postle, McMahon, Ashton, Meredith, & de Zubicaray, 2008). However, the process that leads to formation of associations between novel auditory action words and corresponding motor programs remains essentially unknown.

The majority of previous studies relied on real words with pre-existing semantic representations. Few studies that focused on word acquisition usually examined brain responses to novel linguistic stimuli before and/or after learning (Kelly, McDevitt, & Esch, 2009; Razorenova et al., 2020). In the current study, for the first time, we attempted to investigate the neural changes that occur during the learning process and contribute to formation of an association between a novel pseudoword and a motor action.

We focused on neural oscillations in the beta (β) frequency band (15–25 Hz). Although β-oscillations are ubiquitous across the cortex, their relation to the sensory-motor system has been established most reliably. Sensory-motor β-rhythm is generated in cortico-basal gangliathalamo-cortical loops and robustly modulated by motor performance (Gilbertson et al., 2005). During movement planning and execution, β-oscillations in the sensory-motor cortex are suppressed (β event-related desynchronization (β-ERD)), which indicates the state of active processing (Grent-’t-Jong, Oostenveld, Jensen, Medendorp, & Praamstra, 2013; Heinrichs-Graham et al., 2014; Tzagarakis, Ince, Leuthold, & Pellizzer, 2010). After movement termination, β-power increases, transiently exceeding the pre-movement level (post-movement β event-related synchronization (β-ERS), or β-rebound) (Cheyne, 2013; Pfurtscheller & Lopes da Silva, 1999). Crucially for our study, there is growing evidence that the post-movement β-ERS is implicated in motor learning.

In the studies of motor adaptation, the β-ERS over sensory-motor areas was negatively correlated with movement errors: improved performance after visual-motor training was accompanied by the stronger post-movement β-ERS (Tan, Jenkinson, & Brown, 2014; Tan, Wade, & Brown, 2016; Torrecillos, Alayrangues, Kilavik, & Malfait, 2015). Refinement of hand movements kinematics through repetition of a visual-motor task was associated with an increase in the sensory-motor β-ERS in the hemisphere contralateral to the moving hand (Moisello et al., 2015). Furthermore, the magnitude of this increase predicted retention of the acquired kinematic skill on the following day, so that the higher the increase the greater the retention (Nelson et al., 2017). It was suggested that the sensory-motor β-ERS contributes to learning of motor skills by facilitating the mechanisms of memory formation.

Given the role of the sensory-motor β-ERS in the acquisition of new motor skills, we proposed that a large-scale β-ERS, which comprises cortical regions outside the sensorymotor system, can play a similar role in associative auditory-motor learning. This assumption is supported by two following considerations. First, β-synchronization in the prefrontal cortex and other higher-tier cortical areas was implicated in the maintenance of the existing task rules that govern upcoming responses (the “cognitive set”) in the memory buffer (Engel & Fries, 2010; Spitzer & Haegens, 2017). For example, in experiments using local field potentials (LFP) recorded from the prefrontal cortex in animals, elevation of β-power was repeatedly observed during the delay phase of working memory tasks, during which subjects had to choose a matching stimulus (e.g., Kornblith, Buschman, & Miller, 2016; for review, see Miller, Lundqvist, & Bastos, 2018). But note that EEG/MEG data in humans are rather inconsistent (cf., Deiber et al., 2007; Palva, Kulashekhar, Hämäläinen, & Palva, 2011; for review, see Pavlov & Kotchoubey, 2022). The second consideration relates to the proposed role of β-oscillations in establishing functional neural ensembles distributed over remote cortical and subcortical areas (Antzoulatos & Miller, 2016; Kopell, Whittington, & Kramer, 2011). Such large-scale β-synchronization usually emerges after implementation of a correct decision and it is believed to be engaged in maintaining the “cognitive set” in an active state (status quo -Engel and Fries, 2010), and/or in facilitating synaptic plasticity during learning (Brincat & Miller, 2015).

To test our hypothesis, we analyzed learning-induced changes in β-band modulations using the whole-head magnetoencephalography (MEG) while our participants attempted to acquire associations between the auditorily presented pseudowords and motor responses made by their body extremities under a trial-and-error procedure. We compared the β-power modulations before, during, and after the movements at two stages of learning: (i) the early stage of learning (ESL), when the pseudoword-movement associations had not been acquired yet, and the decisions which particular movement to perform were made rather randomly, and (ii) at the advanced stage of learning (ASL), when each pseudoword cue became associated with the movement of the certain extremity.

Since the associative learning task we used involved motor responses (our participants used one of four limbs to make their choices), there was a possibility that some kind of purely motor learning occurred alongside the associative learning. Distinguishing between the changes in β-modulations within and outside the sensory-motor system was, therefore, central to our study. To address this question, we exploited the well-known contralaterality principle in modulations of β-oscillations originating from cortical representations of the right and left hand (Jurkiewicz, Gaetz, Bostan, & Cheyne, 2006). By separating learning-induced changes on such “effector-specific” sensory-motor β-oscillations and on large-scale β-oscillations that, as we expected, should be common for all four effectors, we hoped to gain insight into the role of β-oscillations in associative learning. The critical prediction was that associative learning should enhance the post-movement β-ERS, with β-band synchronization spreading beyond the sensory-motor cortex and bearing common spatial-temporal patterns regardless of the body extremity used to execute the motor response.

## 2. Materials & methods

### 2.1 Participants

Twenty-four volunteers (mean age 24.9 years, range 19-33 years, 9 females) participated in the main experiment of this study. Additionally, twenty-eight participants (mean age 25.7 years, range 20-38 years, 15 females) performed a self-paced movement task. All participants were native Russian speakers with self-reported normal hearing and no history of neurological or psychiatric disorders. All participants were right-handed according to the Edinburgh Handedness Inventory (Oldfield, 1971). The study was approved by the Ethics Committee of the Moscow State University of Psychology and Education. All participants signed the informed consent prior to enrollment into the study.

### 2.2 Stimuli

The parameters of the stimuli and the procedure of their construction were described in detail elsewhere (Razorenova et al., 2020). Briefly, eight two-syllable pseudowords were constructed in compliance with Russian language phonetics and phonotactic constraints (Table 1). During the associative learning procedure, four of them were unambiguously paired with a movement performed by one of the four body extremities, whereas the other four became associated with no motor response. The first syllable was identical for all pseudowords while the second syllable differed between unique pseudowords: the consonant signaled which extremity (right hand, left hand, right foot, or left foot) a subject might prepare to use, while the vowel specified whether to perform a motor response or to refrain from one. The meaning of the vowel in the second syllable was counterbalanced across stimuli (see Table 1).

**Table 1.**
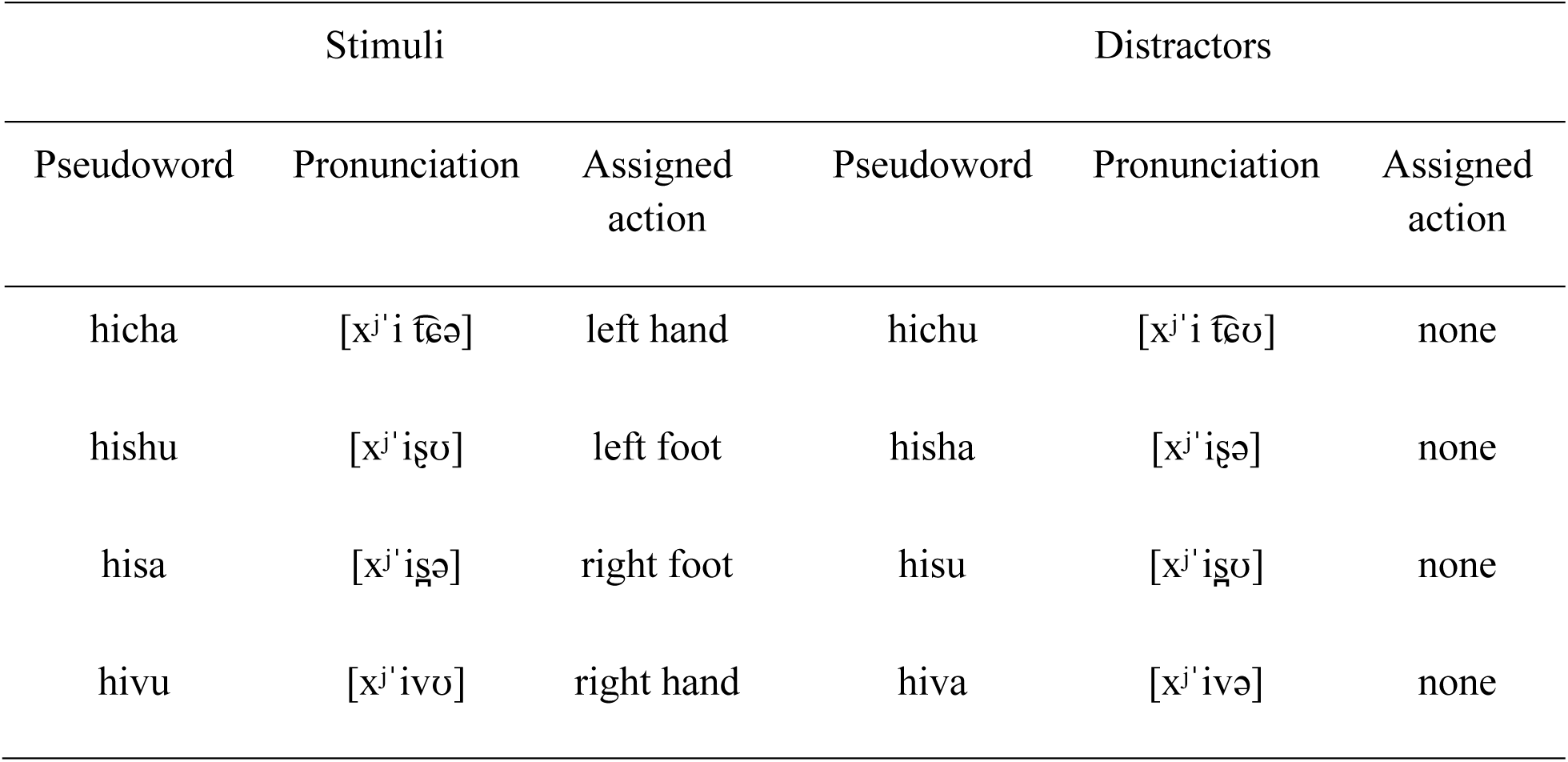
Stimulus-to-response mapping.

The acoustic stimuli were digitally recorded (PCM, 32 bit, 22050 Hz, 1 channel, 352 kbps) by a female native Russian speaker. The pseudowords were constructed by cross-splicing the voice recordings, and the amplitude of the sound recordings was digitally normalized to the maximal power (which corresponded to the stressed vowel), using Adobe Audition CS6.5 software.

Additionally, we used two non-speech auditory stimuli as positive and negative feedback signals. The feedback stimuli were complex frequency-modulated sounds, 400 ms in length each. The signals for positive and negative feedback differed in their frequency range and spectral maxima: for the positive feedback, the frequency range was approximately 400-800 Hz with the spectral maximum increasing in frequency over time; for the negative feedback, the range was 65-100 Hz and the spectral maximum was decreasing during the stimuli.

### 2.3 Design and Procedure

The experiment was implemented using the Presentation 14.4 software (Neurobehavioral Systems, 650 Inc., Albany, CA, USA). The main experiment consisted of four consecutive blocks in the following order: (1) passive listening before learning, (2) early stage of learning (ESL), (3) advanced stage of learning (ASL), and (4) passive listening after learning (Figure 1a). The entire experiment lasted approximately 2 hours.

**Figure 1.**
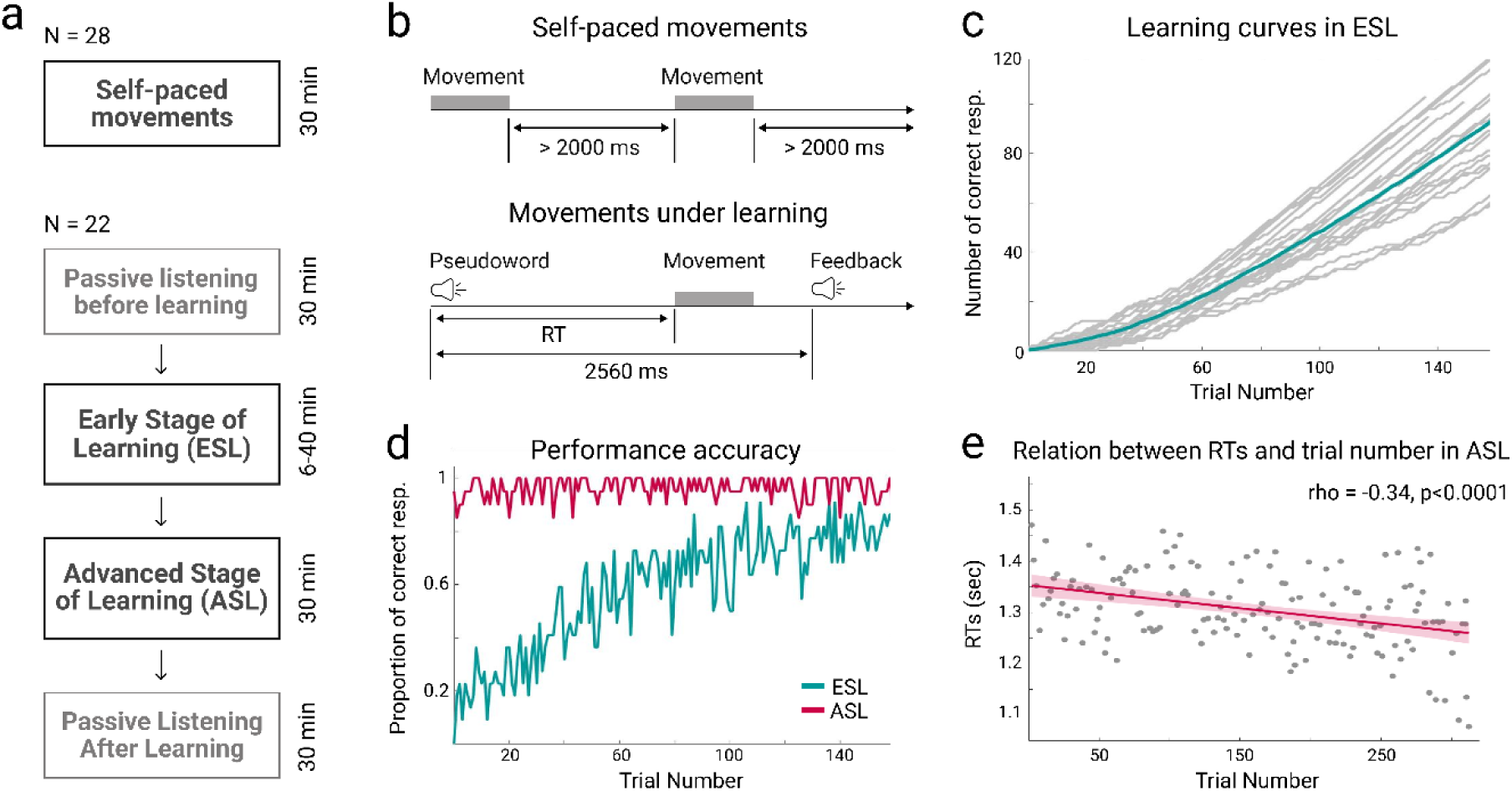
Experimental procedure and behavior. (a) The sequence of experimental blocks. (b) The sequence of events during the self-paced movements and the auditory-motor learning task. (c) Cumulative performance accuracy at the early stage of learning (ESL) over the first 160 trials. The aquamarine line represents the grand average, gray lines -individual data. (d) The proportion of correct responses over time (trial number) in the ESL (aquamarine) and ASL (crimson) conditions. (e) Scatterplot of reaction time as a function of trial number in ASL. The crimson line and semitransparent area around it represent linear regression and .95 confidence intervals respectively. The correlation coefficient and p-value are derived from Spearman rank correlation.

In each block of the main experiment, the participants were presented with the same eight pseudowords (Table 1). The pseudoword stimuli were presented binaurally via plastic ear tubes in a quasi-random order, at 60 dB SPL. The mean duration of the pseudowords was 560 ms.

In the passive listening blocks, participants listened to the pseudowords while watching a silent movie.

During the ESL block, the participants performed a trial-and-error search for the association between each of the presented eight pseudowords and movements of their own body parts. They reacted to each presented pseudoword either by using one of the four body extremities or by making no response; then they received positive and negative feedback signals informing whether their response was correct or erroneous (Figure 1b). The positive feedback was given only if the participant responded in accordance with a predefined pseudoword-to-movement mapping, which was not explicitly revealed to the participants (see Table 1); otherwise, the negative feedback was presented. The ESL block was designed in such a way to meet the requirements of operant learning: the participants tried a variety of new auditory-movement associations and eventually settled upon those that led to positive reinforcement (Neuringer, 2002). The feedback signal was presented at 2560 ms after the stimulus onset (Figure 1b). The next trial began 2400 ms after the feedback stimulus onset with a random jitter of 400 ms. Thus, the total trial length in the active blocks was, on average, 5210 ms.

The length of the ESL block varied depending upon participants’ learning rate: the block was terminated when the participant met a learning criterion or when 480 stimuli were presented, whichever came first. The learning criterion was reached when the participant made at least four correct responses in five consecutive presentations of each of the eight pseudowords. Two out of 24 participants did not reach the learning criterion and thus went through all 480 trials in the ESL block. Their percentage of correct responses in the following ASL block was well within the range of performance of the other 22 participants, therefore, we did not exclude these two participants from further analyses (see Results for details). The average number of stimuli presented within the ESL block was 269 ± 119 (ranging from 74 to 480), with the corresponding duration of the ESL block varying between 6 to 40 min.

The third block (ASL) followed the procedure described above, except that it invariably included 320 trials and lasted approximately 30 min. The fourth block was identical to the first passive block. Short breaks were introduced between all blocks (3 min between the first 3 blocks and 10 minutes before the last one), during which participants were offered to rest while remaining seated in the MEG apparatus.

In a separate group of 28 participants (mean age 25.7 years, range 20-38 years, 15 females), we recorded self-paced movements. During this self-paced block, the participants performed the same movements as those in the main experiment, yet they initiated movements in a self-paced manner, without any external cues. We instructed the subjects to repeat the movement by each body extremity in a series of 30 movements, maintaining an interval of about 2–3 s between the movements. In order to minimize the overlap between the post-movement β-synchronization and preparatory processes for the next movement, we ensured that the interval between any consecutive movements was longer than 2 s. If a participant failed to follow this requirement, she or he was given a pre-recorded warning, and the number of required movements increased correspondingly. Trials with movements separated from the previous ones by less than 2 s were excluded from the further analyzes.

### 2.4 Behavioral responses

Participants performed four movements executed by their body extremities (right hand, left hand, right foot, and left foot). Hand movements were recorded when the participants pressed hand-held buttons (CurrentDesigns, Philadelphia, PA, USA) with their right or left thumb. Foot movements were registered when custom-made pedals were pushed by the toes of the right or left foot. The trajectories of these movements were quite short (< 1 cm for buttons and < 3 cm for pedals), thus, minimizing movement artifacts. The buttons and pedals interrupted a laser light beam delivered via fiber optic cable, and this event was recorded as an onset of the motor response.

The level of performance in the ESL and ASL conditions was evaluated as (i) the mean number of correct responses for four pseudowords that required motor responses and four pseudowords that did not, and (ii) the mean latency of motor responses in the correct trials under the two conditions separately. To characterize the learning curve, we used the cumulative record of conditioned responses as a function of trials (Gallistel, Fairhurst, & Balsam, 2004). The cumulative record is the running sum of the successive behavioral measurements. Changes in the slope of this curve correspond to changes in the level of performance (Figure 1c). Given individual variability in the learning curves, to visualize the group-level course of acquisition under the ESL condition and asymptotic level of performance at the ASL one, we plotted a proportion of subjects, who gave a correct response at each consecutive trial from the first 160 stimulus presentations (Figure 1d). In order to check whether the learning continued in the ASL condition even after the performance became nearly error-free, we computed Spearman rank correlation between the response latency and trial number in the ASL block.

### 2.5 MEG data acquisition

MEG data were recorded inside a magnetically shielded room (AK3b, Vacuumschmelze 699 GmbH, Hanau, Germany), using a dc-SQUID Neuromag VectorView system (ElektaNeuromag, Helsinki, Finland) with 204 planar gradiometers and 102 magnetometers. Prior to MEG recording, the positions of HPI coils and participants’ head shapes (fiducial points and additional points on the head and face) were digitized using the 3D digitizer ‘FASTRAK’ (Polhemus, Colchester, VT, United States). During MEG recording, the subjects’ head positions inside the MEG helmet was assessed every 4 ms. MEG was continuously recorded during each experimental block. Data were sampled at 1000 Hz and bandpass filtered between 0.03–333 Hz with powerline contamination removed by notch-filters centered at 50 and 100 Hz.

The electrooculogram was registered with two pairs of electrodes located above and below the left eye and at the outer canthi of both eyes for the recording of vertical and horizontal eye movements, respectively. A bipolar electromyogram from the right masseter muscle was also recorded for the purpose of muscle artifact detection.

Additionally, the movements of the right and left thumbs were monitored using a 3-axis accelerometer (ADXL335 iMEMS Accelerometer, Analog Devices, Inc., Norwood, MA, USA) at a sampling rate of 1000 Hz. This signal was used later during offline analysis in order to evaluate the fine dynamics of hand movements beyond discretely recorded button-presses.

### 2.6 Data processing

#### 2.6.1 Pre-processing

MEG data were processed off-line. Temporal signal space separation method (tSSS) was applied in order to suppress environmental noise and compensate for head movements (Taulu, Simola, & Kajola, 2005). Then, the data from different blocks and individual subjects were transformed to the standard head position (x = 0 mm; y = 0 mm; z = 45 mm) across all blocks.

MEG data preprocessing was performed using MNE-python software (v0.19) (Gramfort et al., 2013) and custom python scripts. The raw MEG data were downsampled to 300 Hz. The data segments characterized by too low (< 1e−13 T/m) or too high (> 4000e−13 T/m) amplitudes were excluded from the following analysis. Biological (electrocardiographic and vertical electrooculographic) artifacts were removed from the continuous MEG data by means of a semi-automated approach implemented in the MNE-python. We used only gradiometer data for further analyses.

The signals from three accelerometer axes were combined by computing the root mean square at each time point, and then the data were smoothed with a 6-point moving average.

#### 2.6.2 Epoching and time-frequency analysis

In this study, we focused on the ESL and ASL blocks of the main experiment, as well as on the self-paced movements. Given that we were interested in peri-movement β-band modulation, we restricted our analysis to the pseudowords, to which overt motor responses (i.e., button or pedal presses) were executed in both active blocks.

Response-related epochs (comprising 1000 ms before the movement onset and 1400 ms after) were extracted from continuous MEG data. Additionally, for each trial, the time interval from −500 to −100 ms before stimulus onset was taken as a baseline period.

To reject trials contaminated with myographic activity, we filtered MEG data above 70 Hz and calculated the mean of the absolute values across all epochs, at each gradiometer independently. We excluded those epochs during which the maximal amplitude in at least 25% of sensors exceeded 7 standard deviations from the respective means. We also excluded from all the further analyses the epochs with multiple behavioral responses, as preparation and execution of several movements instead of a single movement could modify the neural activity. The trials were then visually inspected to remove the remaining epochs contaminated by bursts of muscle activity. Eventually, 11.7 % of trials were rejected from the ESL block and 11.2% from the ASL block.

For the time-frequency analysis in the sensor space, we used a multitaper power spectrum estimation (Thomson, 2000), implemented in MNE-Python (v 0.19). The sliding time window was 500 ms long with a step size of 25 ms. Time-frequency representations were computed for the 2-40 Hz frequency range in 2 Hz increments, using the ‘tfr_multitaper’ function. Time_bandwidth parameter was set to the default value of 4, resulting in frequency smoothing equal to 8 Hz.

Then, we calculated the power within the a priori selected β-frequency band. In order to prevent an overlap between the effects in the β-frequency range and effects in adjacent frequency bands, we restricted further analyses to a conservative range of 15-25 Hz. Frequencies below 15 Hz were excluded from the analyses because the range of αoscillations in MEG data may extend up to 15 rather than 12-13 Hz (Mierau, Klimesch, & Lefebvre, 2017). We restricted the analysis to frequencies below 25 Hz in order to avoid possible effects in the γ-range (Hashimoto et al., 2017). The resulting 15-25 Hz frequency range was previously used in many studies on the role of β-oscillations in linguistic processing (Klepp, Niccolai, Buccino, Schnitzler, & Biermann-Ruben, 2015; Niccolai et al., 2014; Pavlova et al., 2019; L. Wang et al., 2012; Weiss & Mueller, 2012). Visual inspection of time-frequency plots at MEG sensors showed that the chosen frequency range was strongly affected by our auditory-motor learning task (see Suppl. Figure 1 for a representative sensor) thus encouraging its further analyses.

For each participant in each condition, we added power values for all 2-Hz frequency bands within the 15-25 frequency range at each time point of the selected epochs. To reduce intersubject variability and to normalize β-power changes, the data were baseline-corrected by dividing β-power at each time point by the averaged power computed over the pre-stimulus interval (from −500 to −100 ms relative to stimulus onset). The resulting values were converted to dB by log10-transformation and multiplication by 10. Planar gradiometers were then combined by adding time-frequency data for pairs of channels corresponding to orthogonal sensors at the same location. All analyses at the sensor level were thus performed at 102 combined gradiometers.

We analyzed the movement-related β-power during the self-paced condition using the same procedures as described above. The baseline interval was taken from −1000 to −600 ms before the movement onset.

#### 2.6.3 Analysis of effector-specific effects at the sensor level

In order to identify the effects that the learning task had on sensory-motor β-oscillations, we exploited well-known hemispheric asymmetry in their modulations during left-hand and right-hand movements, i.e., the contralateral predominance of the β-ERD/ERS in the sensorymotor areas relative to the side of the hand movement (“effector-specific” β-effects).

For each experimental condition (Self-paced movements, ESL and ASL) we extracted 12 trials with left-hand movements and 12 trials with right-hand movements (24 trials per condition in total). The trials were selected from the beginning of the Self-paced and ESL blocks, and from the end of the ASL block. To ensure that changes in β-power between the ESL and ASL conditions were induced solely by learning, we matched the ESL and ASL trials both in terms of stimuli presented and hands used to make the response. Only correct trials were selected for both the ESL and ASL conditions, even though for the ESL block a correct response might be a matter of chance, especially at the beginning of the condition. At this stage, two participants were excluded from further analyses because of insufficient numbers of correct trials in the ESL condition. The final sample included 22 subjects.

We utilized the Self-paced condition as a functional localizer to determine which MEG sensors were preferably suited for capturing “sensory-motor” β-band modulations. We chose the three time intervals of interest (each 450 ms in length) (see Results): (1) “Pre-movement” interval coinciding with movement preparation − from −550 ms to −100 ms before the movement onset; (2) “Movement” interval representing movement execution − between 100 and 550 ms after the movement onset; (3) “Post-movement” interval − from 550 to 1000 ms after the movement onset. Based on the previous literature (e.g., Cheyne, 2013), we identified “sensory-motor” sensors as the ones that displayed the strongest β-ERD during movement preparation. For this purpose, individual minima in the β-ERD values within the Premovement time interval were identified for each of 102 combined gradiometers, for left- and right-hand movements separately. The grand average across participants was visualized at scalp topographic maps, and six combined gradiometers (three for each hand) displaying maximal suppression of β-power were selected.

To check whether the gradiometers chosen in this way capture contralaterality effect both in the β-ERD and in the maximal post-movement β-ERS, for the selected sensors we evaluated individually defined maxima in the β-power during the Post-movement time interval. The respective maxima of the pre-movement β-ERD or the post-movement β-ERS were averaged across the selected sensors separately for the left- and right-hand movements and for contra- and ipsilateral hemispheres. The resulted values were subjected to the β-ERD and β-ERS rmANOVAs with two factors: Hand (left- or right-hand movements) and Contralaterality (contra- and ipsilateral relative to the performed movement). For visualization purposes, we also plotted time courses of β-power change averaged across the three “sensory-motor” sensors during the contralateral hand movements. In order to estimate the time intervals of the significant β-ERD and β-ERS, we contrasted the normalized β-power value at each time point with a reference value of zero using a single-sample t-test. The statistical results were FDR-corrected for the number of time points (n = 96).

We used the same procedure to probe for contralateral bias in effector-specific β-power modulations at the selected «sensory-motor» gradiometers during hand movements performed in associative learning conditions. The individual minimal and maximal β-values at the sensors of interest were subjected to rmANOVAs with 3 factors: Hand, Contralaterality, and Condition (ESL and ASL). This analysis was complemented by point-by-point comparison of the learning-induced changes in the time courses of the sensorymotor β-power for the right- and left-hand trials by means of two-tailed paired t-test. We also compared baseline-normalized β-power in each point of those time courses to a reference value of zero using a single-sample t-test. All statistical results were FDR-corrected for the number of time points (n = 96).

As long as the behavioral task required multiple motor responses in both ESL and ASL conditions, we anticipated that some type of motor learning could take place during the experiment. To check this possibility, we contrasted the accelerometer signals recorded during the hand movements under the ESL and ASL conditions using paired t-test. The differences in movement duration between the two conditions (see Results for details) promoted the following supplemental analysis.

We repeated the rmANOVAs described above in a subset of participants whose accelerometer signal did not differ between the conditions. We selected such participants using the following procedure. First, we calculated the mean accelerometer signal across left and right-hand movements within a broad time window from 100 up to 1000 ms after the movement onset in each condition separately. Then, we ranged the participants according to the magnitude of ESL minus ASL difference in the averaged accelerometer signal. We excluded participants with the highest rank one by one until the difference in the accelerometer signal was no longer significant according to the single-sample t-test. As a result, 13 participants were included in the subsample. Their data were subjected to the same analyses as those described above.

#### 2.6.4 Analysis of the large-scale effects at the sensor level

Further analyses of learning-induced changes in β-power modulations were focused on the large-scale effects. We expected that β-power modulations resulting from cognitive aspects of associative learning would be independent of the body extremity used for a response and would comprise a large part of the sensor array. We tested both assumptions using only responses performed by hands.

To this end, we evaluated the whole-head topography of β-power changes in the three response-locked time intervals that were analyzed separately for the left- and right-hand trials. Normalized β-power at each combined gradiometer for each participant and hand movement (left or right hand) was averaged over time points within Pre-movement, Movement, Post-movement intervals. To evaluate the degree and direction of the β-power changes relative to the baseline, we compared the β-power values in each interval and under each condition to a reference value of zero using a single-sample t-test. The statistical results were FDR-corrected for 102 combined gradiometers, three time-intervals and 2 hand movement (612 comparisons in total), and represented as topographic plots. Then, under the null hypothesis of no differences in the observed large-scale β-modulations between left- and right-hand movements in either experimental condition, the averaged β-power values at each sensor and time interval during left-hand movements were subtracted from the ones during the right-hand movements and contrasted with a reference value of zero by means of the single-sample t-test. The analysis was supplemented by the sequence of topographical maps of β-power modulations at successive 200 ms intervals separately for movements performed by the right and left hands, as well as topographical maps of their differences.

When the null hypothesis was confirmed, we proceeded with the analysis of effector-nonspecific large-scale β-modulations, while collapsing the trials with movements of all four body extremities for further comparisons. Within each condition, we took into analysis an equal number of correct trials for each extremity (namely, seven for each extremity, thus, producing 28 epochs in total). For the ESL condition, 28 epochs were extracted from the beginning of the ESL block, while, for the ASL condition, 28 trials were taken from the end of the respective block. Then, we calculated mean β-power changes in each time interval in both conditions following the same steps of analysis as described above for the trials containing only the hand movements. The results of the analysis were displayed as topographical maps.

To explore the learning-induced changes in the temporal dynamics of large-scale β-modulations, the β-power change values from combined gradiometers of the left half of the sensor array and from gradiometers of the right half were averaged for each time point for the ESL and ASL conditions separately. The resulting time courses were contrasted between the two stages of learning by means of paired t-tests (FDR-corrected for the number of time points). In addition, we averaged β-change values across Movement and Post-movement time intervals (100-1000 ms) for each side of the sensor array and each condition separately and subjected them into rmANOVAs with two factors (Hemisphere, Condition).

The further analyses were performed using the same set of correct trials containing movements of all four body extremities under the ESL and ASL conditions.

At the final step of the sensor-level analysis, we examined whether the increase in the normalized β-power in the ASL condition at the remote sensors occurred in the same or different trials. To do so, we selected one sensor in the left frontal, one in midline posterior, and one in the right temporal parts of the sensor array (see Results for details) and averaged the β-power over the Movement and Post-movement intervals merged together (100-1000 ms after the movement onset) in each trial in each of the three selected sensors. The resulting β-power values were subjected to regression analyses using linear mixed-effects models (Bagiella, Sloan, & Heitjan, 2000).

We built three models where the dependent variable was the value of β-power change on one sensor and the independent variable was the β-power change on the other. The mixed-effects models were created with the ‘lmer()’ function available in the lme4 package for R (Bates, 2010). Differences in β-power between subjects (22 levels) and between the limbs the participants responded with (4 levels) were attributed to random effects. We assumed that the subjects may have a different magnitude of the task-induced β-power change and different movements may trigger different changes in β-power, that is why we used separate intercepts for subject and limb taken as random factors in the model. The models were represented in the following way:

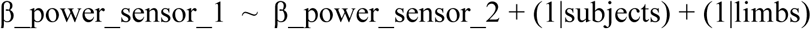

For visualization purposes, we plotted scatterplots that depict the relationship between the β-power change on each pair of the sensors.

#### 2.6.5 Source localization of the large-scale effects

Structural MR scans (voxel size 1 × 1 × 1 mm) were performed on a 1.5T Philips Intera system. The T1 images were processed using the default ‘recon-all’ reconstruction algorithm (Fischl et al., 2002), implemented in FreeSurfer 4.3 software (Martinos Center for Biomedical Imaging, Charlestown, MA). The results of this reconstruction were used to create the source model as described below.

Single layer BEM models were created for every subject based on the anatomical MRI using the FreeSurfer watershed algorithm (Ségonne et al., 2004). MEG data were co-registered with each subject’s structural MRI using the ‘mne_analyze’ tool (MNE-software (Gramfort et al., 2014)). Next, the forward model was estimated using the surface-based source space with 10,242 vertices per hemisphere. Noise covariance matrices were estimated from the baseline period (−500 – −100 ms relative to stimulus onset). To reconstruct cortical sources of the power changes within 15-25 Hz frequency band, we applied a dynamic statistical parametric mapping (dSPM) localization method (Dale et al., 2000) in conjunction with an eight-cycle complex Morlet wavelet using MNE toolbox default options (function ‘source_band_induced_power’). Vertices within a non-cortical medial wall area (Glasser et al., 2016) were excluded from further consideration. Then, to compute the time courses of the induced β-power changes, the data were baseline-corrected by dividing the β-power at each time point by the averaged power computed over the baseline interval from −500 to −100 ms relative to stimulus onset. The resulting values were converted to dB by log10-transformation and multiplication by 10.

To evaluate the cortical topography of β-power modulations during initial and advanced learning stages, we averaged the data in each vertex within each of the three time intervals of interest (Pre-movement, Movement, and Post-movement) in the ESL and ASL conditions separately. The resulting log-transformed baseline corrected values were contrasted with the zero using the single-sample t-test. The p-values in each condition were FDR-corrected for the number of vertices and the number of time intervals (n = 10242*2*3 = 61452).

To unravel the temporal dynamic of the β-ERS at the source level, we chose five cortical labels from Destrieux Atlas (‘aparc.a2009s’ in Freesurfer; Destrieux, Fischl, Dale, & Halgren, 2010) which overlapped with the peaks of the statistical map for the early β-ERS in the left hemisphere during the ASL condition as was discovered at the previous step of source level analysis (see Results for details). We also considered the symmetrical labels from the right hemisphere. Within each of the ten selected labels, we found ten most significant vertices and averaged values of β-power change across them at every time point during the epoch. The resulting time courses were contrasted against the pre-stimulus baseline by means of a single-sample t-test. All p-values were FDR-corrected for the number of time points.

#### 2.6.6 Relationship between β-power and behavior

In order to examine the dependence between the β-power and the participants’ behavior, we used LMEM analysis. In each trial, we used minimal β-values during the Pre-movement interval and maximal β-values during the Post-movement interval in the ESL and ASL conditions as a dependent variable. The independent variable was response time (RT) in corresponding trials. The mixed-effects models were built with the ‘lmer()’ function available in the lme4 package for R (Bates, 2010). Differences in β-power between subjects (22 levels) and between the limbs the participants responded with (4 levels) were attributed to random effects. The full model was represented in the following way:

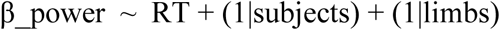

Then, we tested the model at every combined planar gradiometer, with Pre-movement and Post-movement intervals and each condition treated separately, the p-values were FDR-corrected for the number of sensors in each time interval in each condition (n = 102 sensors). The regression coefficients were plotted on topographical maps.

For visualization purposes, in each of the four cases (two intervals and two conditions), we chose the most significant sensor (if any sensors were significant) and drew a scatterplot that depicts the relationship between the β-values at this sensor and RT at a single-trial level. For this plot, in order to get rid of the differences between limb movements (e.g., the participants took consistently more time to respond with legs than with hands), we converted individual RT into z-scores for each limb separately.

## 3. Results

### 3.1 Behavioral results

All, but two, participants reached the learning criterion in the ESL condition, with 269 ± 119 (mean ± standard deviation) stimuli presented within the block. Two participants, who did not reach the learning criterion and thus went through all 480 trials, performed successfully in the ASL condition, with a percentage of correct responses well within the range of performance of the other 22 participants (94% and 98% vs. the range of 86-99%); therefore, we did not exclude them from the further analyses. Two participants reached the learning criterion before they had performed the number of trials sufficient for statistical analyses (at least seven trials for every moving body extremity, see Method); thus, we excluded them from the further analyses. The final sample included 22 participants.

The ESL and ASL conditions significantly differed in the total number of correct responses: the mean percentage of correct responses was 68 ± 6.7% (ranging from 57% to 80%) in the ESL and increased up to 96 ± 3.6% (86% −99%) by the ASL condition (t(21) = −19.192, p < 0.0001). In the course of learning, mean response time became significantly shorter in the ASL condition compared with the ESL condition (1406 ± 111 vs 1322 ± 116 ms, t(21) = 3.531, p=0.002).

These data reveal that, when the participants acquired the contingency between the stimuli and corresponding body extremities, their decision to execute a cued motor response became easier. Notably, even within the ASL condition taken separately, there was a significant negative correlation between the successive trial number and the response latency (rho = −0.34; p<0.0001) thus indicating progressive shortening of mean motor response latency throughout this block (Figure 1e). This suggests that further strengthening of auditory cue-motor association occurred even during the advanced learning stage when the participants already performed the task with high accuracy.

### 3.2 Effector-specific β-power modulations

To separate the impact of auditory-motor learning specifically associated with sensory-motor processing from more general effects of rule acquisition, we exploited the well-established contralateral dominance in β-power modulation induced by execution of hand movements in the hand area of the human sensory-motor cortex.

To identify the MEG sensors preferably suited for capturing effector-specific sensory-motor β-modulations, we used self-paced movements executed by the left and right hands separately as a localizer task. Figure 2a depicts the topographic maps of induced β-power changes averaged within each of the three 450 ms periods – before, during, and after movement – for each moving hand separately, as well as the differential topographic maps (right- versus left-hand movements). In accord with the numerous prior evidence (e.g., Pfurtscheller & Lopes da Silva, 1999; Tzagarakis et al., 2010), movement preparation induced significant β-power reduction (β-ERD) in the combined gradiometers overlying sensory-motor hand areas, that, although weakening, sustained throughout the movement execution period (Figure 2a,d). After movement cessation, a β-power enhancement slightly above the baseline level (β-rebound) was significant for the right-hand movements, although a similar non-significant tendency could be observed for the left-hand movements (Figure 2a,d). The expected contralateral dominance was clearly evident during the Pre-movement interval, with the significant β-ERD mostly confined to the hemisphere contralateral to the moving hand.

**Figure 2.**
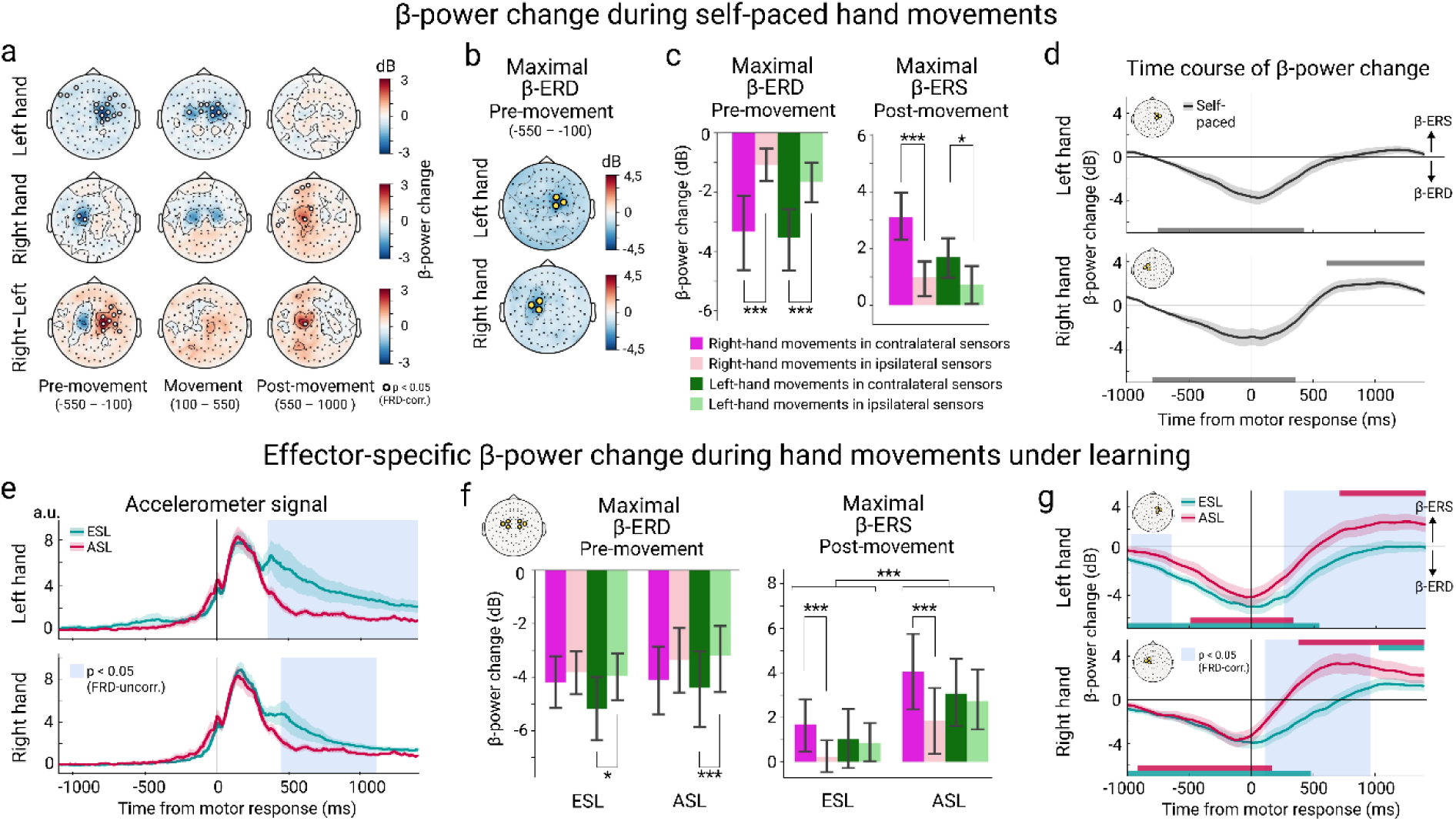
Effector-specific movement-related β-power change during hand movements in self-paced (a-d) and learning conditions (e-g). (a) Topographic maps of β-power change (15-25 Hz) during self-paced hand movements. Baseline normalized β-power is averaged within each of the three time intervals: before, during, and after movement. The upper and medium rows represent movements of the left and right hands respectively, and the bottom row depicts the contrast between the left- and right-hand movements. Open circles indicate sensors with significant β-power changes relative to the pre-stimulus baseline (the two upper rows) or significant difference in β-power between the left and right movements (the bottom row) (p < 0.05, FDR-corr.). Here and hereafter, blue and red hues mark the β-ERD and β-ERS respectively. (b) Topographic maps of the maximal β-ERD values during the Pre-movement interval. The yellow circles indicate the «sensory-motor» sensors with the strongest β-ERD for movement of the contralateral hand. (c) Bar graphs represent the maximal pre-movement β-ERD (left panel) and post-movement β-ERS (right panel) in the «sensory-motor» sensors at the left and right hemisphere during contra- and ipsilateral hand movements. Here and hereafter, whiskers indicate the 95% confidence intervals. Asterisks denote the level of significance: *p<0.05, **p<0.01, ***p<0.001. (d) Time courses of β-power change in the “sensory-motor” sensors for contralateral movements. A vertical line corresponds to a motor response onset. The shaded area around a time course represents the standard error of the mean (SEM). Here and hereafter, horizontal bars below and above a time course mark time intervals with the significant β-ERD and β-ERS respectively (p < 0.05, FDR-corr. for a number of time points). (e) Grand average of accelerometer signal during left- and right-hand movements in ESL and ASL. Here and hereafter, aquamarine color denotes ESL, crimson color – ASL condition. Blue shaded area marks the time interval with the significant ESL-ASL difference (p < 0.05, uncorr.). (f) The maximal pre-movement β-ERD and post-movement β-ERS in the “sensory-motor” sensors during contra- and ipsilateral hand movements in the ESL and ASL. All designations as in Figure 2c. (g) Time courses of β-power in the “sensory-motor” sensors for contralateral movements in ESL and ASL. Blue shaded area marks time interval with a significant ESL-ASL difference (p < 0.05, FDR-corr. for a number of time points).

Location of the three combined gradiometers capturing the strongest individual β-ERD during the Pre-movement interval was exactly symmetrical for the left- and right-hand movements (Figure 2b). For the following analysis, the minimal β-values from the PreMovement interval (β-ERD) and the maximal β-values from the Post-movement interval (β-ERS) were averaged over the respective sensors and subjected to rmANOVAs with factors Hand (right/left) and Contralaterality (contralateral/ipsilateral hemisphere). The contralaterality effect was significant in both analyses (β-ERD: F(1, 21) = 35.8, p < 0.0001, η^2^ = 0.14; β-ERS: F(1, 21) = 23.5, p = 0.0001, η^2^ = 0.14; see also Figure 2c for the planned comparisons). This confirmed that for the self-paced movements, the β-ERD and β-ERS in the selected triplets were always strongest in the contralateral hemisphere, thus complying with the distinguishing feature of the effector-specific sensory-motor β-oscillations. Therefore, these six combined gradiometers (three for each hemisphere) were taken for further analysis.

Then, we explored (i) whether a similar contralateral predominance could be observed for the effector-specific β-ERD and β-ERS induced by the auditory cue-triggered hand movements in the associative learning task, and (ii) whether the learning progress changed the strength of movement-related β-modulations.

To this end, the individual minima of β-power in the Pre-Movement interval (β-ERD) and maxima in the Post-movement interval (β-ERS) were averaged over each of the gradiometer triplets for the ESL and ASL conditions separately and subjected to rmANOVA analyses. The factors were Hand (right/left), Contralaterality (contra-/ipsilateral hemisphere) and Learning stage (ESL/ASL). For both the β-ERD and β-ERS analyses, the main effect of Contralaterality was significant (β-ERD: F(1, 21) = 15.03, p = 0.0009, partial η^2^ = 0.41, and β-ERS: F(1, 21) = 17.38, p = 0.0004, partial η^2^ = 0.45), thus indicating that sensory-motor β-modulations during learning task were greater in the sensory-motor areas of the hemisphere contralateral to a moving hand (see Figure 2f for the planned comparisons). Notably, during the task, the contralaterality effect for the post-movement β-ERS (but not for the pre-movement β-ERD), was stronger for the right-hand than for the left-hand movements (Hand × Contralaterality interaction (F(1, 21) = 9.94, p = 0.0048, partial η^2^ = 0.32; see Figure 2f). The transition from the early to advanced learning stage was associated with increased strength of the β-ERS (Condition effect: F(1, 21) = 16.16, p = 0.0006, partial η^2^ = 0.44), while it did not significantly affect the β-ERD (F(1, 21) = 0.87, p = 0.36). No other factors or interactions were significant.

Thus, auditory-motor learning led to enhanced effector-specific β-synchronization occurring after the movement termination. Yet, it is conceivable that, in the course of the auditory-motor learning, some parameters of movement execution could change (e.g., the participants may decrease the duration of the button press or reduced the number of subthreshold attempts to perform corrective movements, etc.); that, in turn, could affect the dynamics of βoscillations (e.g., Pakenham et al., 2020). To check whether this effect was present in our data, we compared the accelerometer signals for movements executed by right or left hands in the two conditions, and we did observe a tendency to shorten movement duration as learning advanced from the ESL to ASL conditions (a relative decrease of grand average accelerometer signal in the 450-1100 ms after movement onset; Figure 2e) (p < .05, uncorr.). This suggests that the weaker β-ERS in the Post-movement interval in the ESL as compared with ASL might be a mere consequence of relatively more protracted movements extending into the selected post-movement interval at the beginning of learning.

To test whether movement duration could affect MEG data, we repeated the rmANOVA on the β-ERS maxima in a subset of 13 participants whose averaged accelerometer signals within the 100-1000 ms time interval after the movement onset was closely matched between the ESL and ASL conditions (Suppl. Figure 2; see Methods for details). The main effect of Condition remained significant (F(1, 21) = 9.31, p = 0.01, partial η^2^ = 0.44). This finding strongly suggested that the observed learning-related enhancement in the effector-specific post-movement β-ERS could not be not solely explained by motor training.

### 3.3 Large-scale β-power modulations

Along with the β-band modulation in the sensory-motor cortex, we also expected learning-related effects in the β-frequency range outside the sensors overlying the sensory-motor cortex. For this purpose, we analyzed the ESL-ASL difference across the whole sensor array, thus avoiding confounds bound to the selection of the pre-defined sensory-motor ROI. Figure 3a displays the topographic maps of β-modulation for the left- and right-hand movements in the ESL and ASL conditions separately. When the movements were executed in the context of the learning task, they were accompanied by the large-scale β-power modulations, which were not restricted to the “sensory-motor” regions. Rather, the sensors that captured the significant pre-movement β-ERD under the ESL and the post-movement β-ERS under the ASL conditions comprised a large portion of the entire sensor array, with a visible left-hemispheric bias in the latter case. Notably, under the ASL condition, the left non-sensorymotor sensors demonstrated the significant β-ERS during the movement execution period itself, i.e., long before the movement was terminated. These large-scale pre-movement β-ERD and post-movement β-ERS were not effector-specific, since they did not significantly differ between the left- and right-hand movements either under the ESL or ASL condition (Figure 3a, bottom row). Moreover, when we collapsed data across the hand and foot movements (Figure 3b), the topographical maps of β-power modulations for all four movements pooled together essentially reproduced the general pattern observed during the left and right hands movements. All further analyses were performed on the data pooled across four body extremities.

**Figure 3.**
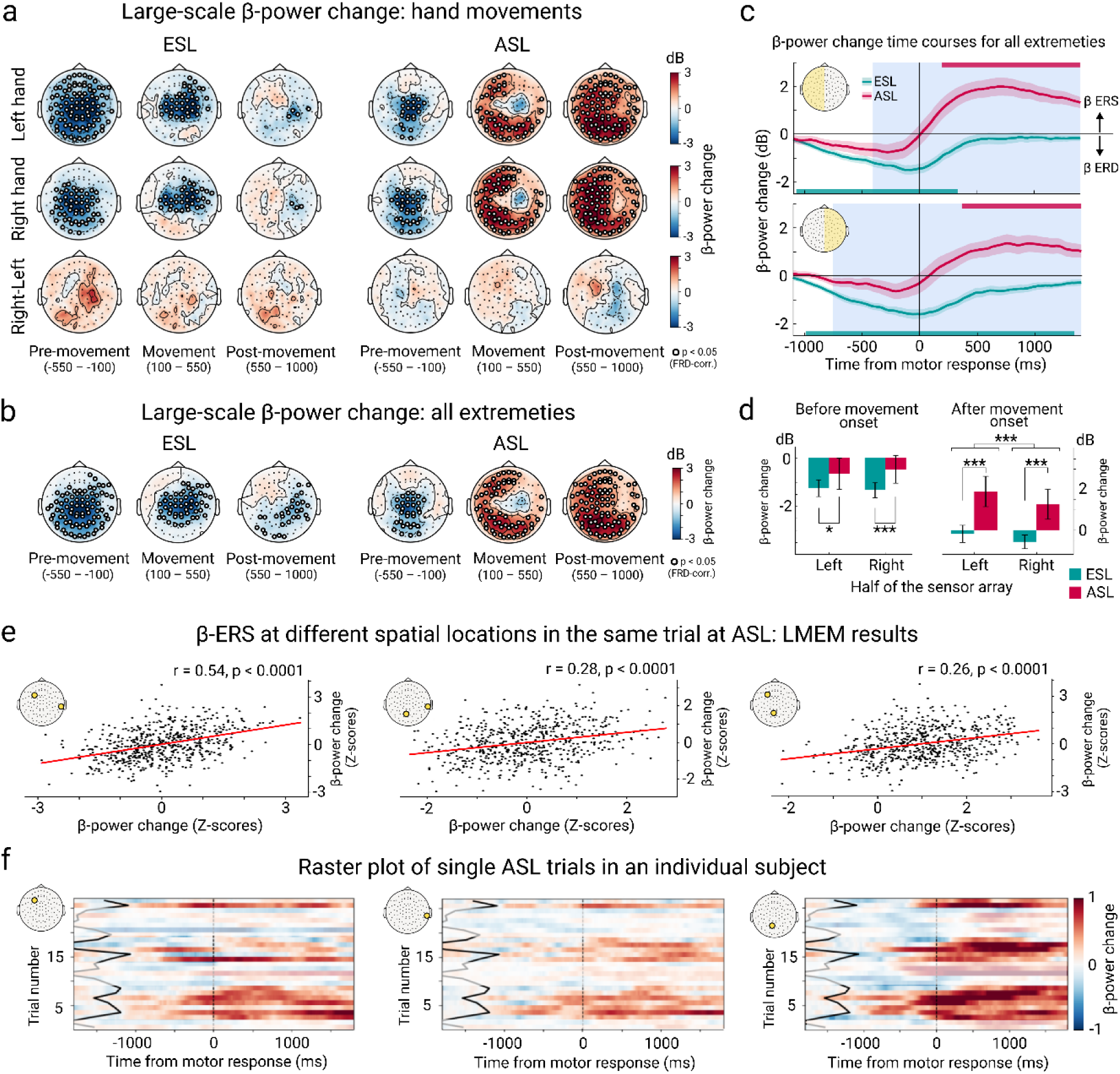
Large-scale β-power change related to movements at early and advanced stages of learning (ESL and ASL). (a) Topographic maps of β-power change related to the hand movements. (b-f) β-power change for movements of the left and right hand, left and right foot pooled together: (b) Topographic maps; (c-d) β-power change averaged within the left and right halves of the sensor array: movement-related time courses (c) and mean values within two time intervals: before (−550 – −100 ms) and after (100-1000 ms) the movement onset (d); (e) The relationships between the ASL β-ERS at different spatial locations at the single-trial level. The scatterplots are for display purposes only, regression coefficients and p-values in the upper right corner are derived from the linear mixed effect models (LMEM). The × and y axes represent the individual z-scores of the post-movement-onset β-ERS in each trial for the selected pair of MEG channels (their location is shown on the insert head); (f) Raster plots of ASL β-power change as a function of the trial number in a representative subject. Trials are sorted in the order of their presentation. The horizontal raster lines represent single trials in a subject’s dataset. Blue color -β-ERD, red -β-ERS. The yellow dot on the scalp map (top left) indicates the position of the channel. A vertical dotted line marks the movement onset, the solid angled line connects stimulus onsets in each trial. Note the occurrence of the strong and protracted β-ERS in all three spatial locations in the same trials.

First, to roughly characterize the large-scale β-modulations emerging in the context of learning, and to explore their putative changes with learning progress, we computed the time courses of the baseline-corrected β-power values averaged over the fifty-one combined gradiometers constituting the left and right parts of the sensor array for each learning condition separately (Figure 3c). Under the ESL condition, the right-hemispheric β-power time courses revealed the protracted β-ERD, which was sustained until the end of the Post-movement period (1000 ms after movement onset), while in the left hemisphere the β-ERD returned to the baseline values already during movement execution (∼250 ms after the movement onset). No significant the large-scale β-ERS was present in both hemispheres. In sharp contrast to the ESL, in the ASL condition, the significant β-ERD in both hemispheres returned to the baseline levels by the end of the Pre-movement period. Shortly after movement onset (∼200 ms relative to a movement onset), it was substituted by the highly significant β-ERS in the left hemisphere, which also emerged with ∼50 ms delay in the right hemisphere, and then sustained in both hemispheres until the end of the Post-movement period.

To verify and extend the emerging picture of learning impact on the large-scale β-modulations, we performed the rmANOVAs for the average left and right-hemispheric β-power values that were extracted from the two successive time intervals relative to the movement onset. Given that beginning of the movement, rather than its cessation, represented a turning point when the large-scale β-ERD was replaced by the β-ERS, the mean β-power in left and right hemispheres in each learning condition were averaged within the two time intervals, one of which preceded and the other one followed the movement onset (−550 − −100 ms, and 100 − 1000 ms relative to a movement onset). The rmANOVA with factors Condition (ESL/ASL) and Hemisphere (left/right) revealed that for the pre-movement interval the only significant factor was Condition (F(1, 21) =10.35; p = 0.004; partial ɳ^2^ =0.33). This was explained by significant learning-induced attenuation of the large-scale β-ERD strength occurring during movement preparation (Figure 3c,d). For the post-movement onset interval, on the other hand, the significant effect of Condition (F(1, 21) =36.37, p<0.0001; partial ɳ^2^ =0.63) was mainly due to the emergence of the highly pronounced largescale β-ERS in the ASL, which was absent in the ESL condition. The greater strength of the learning-induced β-ERS in the left than in the right hemisphere was reflected in the significant main effect of Hemisphere (F(1, 21) =, p <0.0001; partial ɳ^2^ =0.60; Figure 3d).

To sum up, early in the auditory-motor learning, the cue-triggered movements were mainly accompanied by the large-scale β-ERD that was sustained even after the movement cessation. As learning advanced, the β-ERD was greatly attenuated and replaced by the long-lasting widespread β-ERS, which evolved around the time of the movement onset and had a significant left-hemispheric predominance. This suggests that rather than being triggered by movement termination (as it was observed for the effector-specific β-ERS), the rise of the large-scale β-ERS was linked to the end of the decision-making process regarding movement selection.

The large-scale β-ERS was unevenly distributed over the sensor array (Figure 3b). Spatial clusters of combined gradiometers that showed maximal increases in synchrony in the β-band before movement termination were located over the left frontal and posterior parietal regions, i.e., far away from each other, and were separated by locations with the much less prominent β-ERS. This substantially reduced the possibility that the widespread effect could be produced by power leakage between the frontal and parietal β-ERS clusters. On the other hand, since we used the across-subject and across-trial averaging approach, it remained unclear whether the β-band activity in the distant clusters occurred in the same or different trials. To examine this more closely, we performed a single-trial power-power correlation analysis between three distant gradiometer channels located over the left frontal, midline posterior parietal, and right temporal areas using the LMEM approach (Figure 3e). The results showed that in a single ASL trial, increased β-power at one spatial location cooccurred with the increased β-power at the rather distant location (β coefficients varied from 0.54 to 0.26; all ps<0.0001; see also Figure 3f for a representative subject). Thus, the singletrial β-ERS was modulated in a coordinated manner across different spatial locations.

To assess the neural sources of the learning-induced effects that we have statistically demonstrated at the sensor level, we performed the source localization analysis. For the ESL and ASL conditions separately, we localized the cortical sources of the mean β-power modulation within each of the same time intervals – before, during, and after the movements – that were used for the sensor-level analysis. Consistent with the scalp topographies in Figure 3b, early in the learning, the pre-movement β-ERD predominantly occurred in the regions of the left and right parietal lobes at their lateral and medial surface, but also at the posterior part of medial and lateral frontal regions bilaterally (see Figure 4a, upper row). The β-ERD at this initial learning stage continued throughout the entire time interval up to the end of the Post-movement period. Under the ESL condition, no significant rise of β-power above the baseline values was detected in any cortical sources. The vertex-wise ESL-ASL contrast confirmed highly significant attenuation in the β-ERD strength after the successful acquisition of the associative rules (ps<0.001, FDR-corrected, Suppl. Figure 4). Learning progress toward the ASL condition was associated with the appearance of the early β-ERS evolving long before the movement termination, which was localized in the dorsolateral, ventrolateral and medial prefrontal regions of the left hemisphere alongside with medial surface of the occipital and temporal lobes (Figure 4a, bottom row). After movement cessation, the learning-induced β-ERS was further enhanced in these regions and spread over the cortical areas of the lateral parietal and temporal lobe. Although the widespread postmovement β-ERS became bilateral but it still remained more extensive in the left hemisphere, again consistent with the sensor-space results presented in Figure 3b. Notably, the source-space analysis indicated involvement of a fairly broad swath of anterior and posterior regions of the cingulate and retrosplenial cortex in the learning-induced β-ERS.

**Figure 4.**
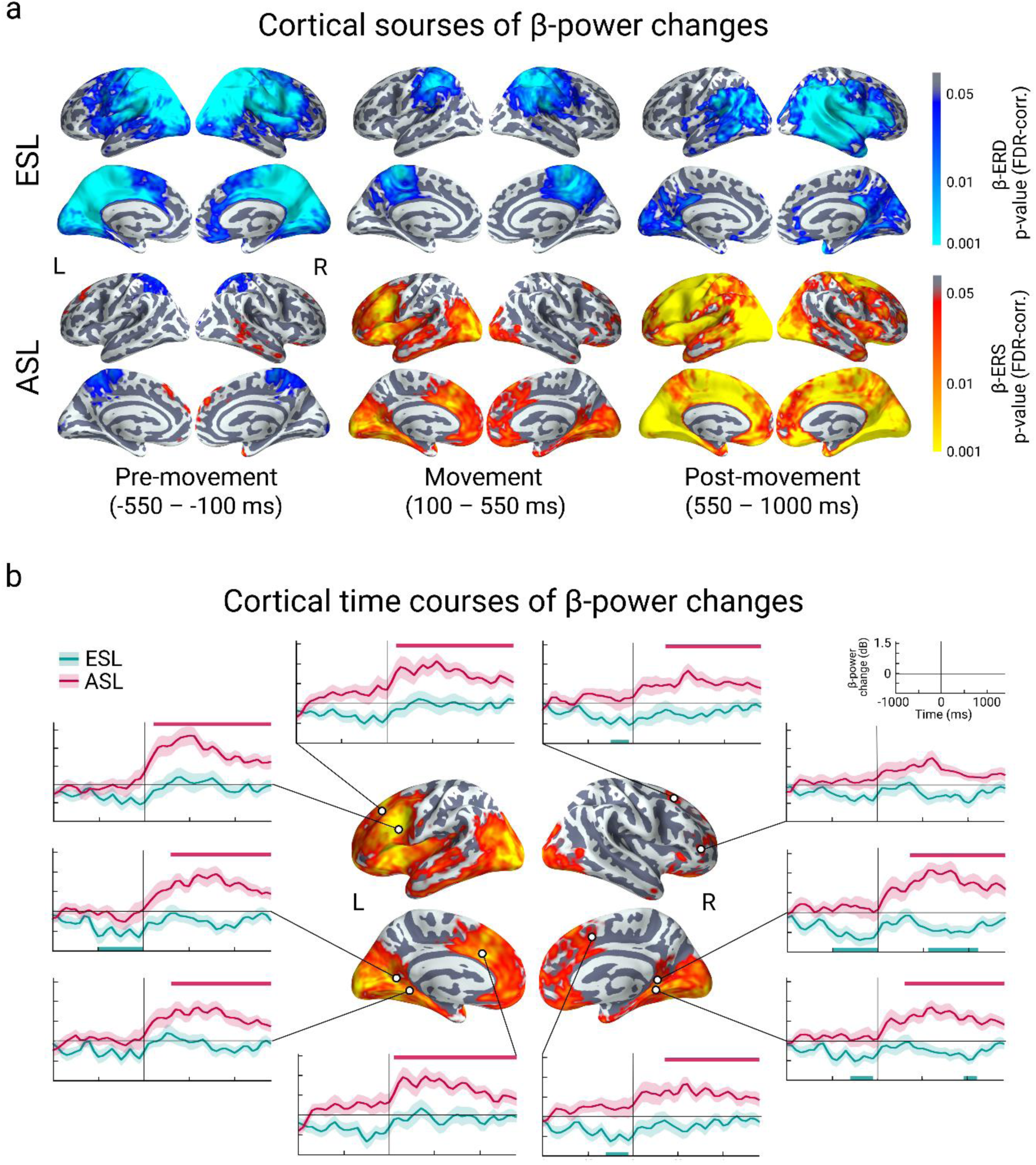
Cortical sources of β-ERD and β-ERS during ESL and ASL. a) Statistical cortical maps of β-power change in successive time windows in ESL and ASL. Blue hues indicate cortical regions with the significant β-ERD and red-yellow hues – with the significant β-ERS (p < .05, FDR-corr.). b) Time courses of β-power change at the source level. The cortical map in the middle displays the areas with the significant β-ERS in the 100-550 ms interval after movement onset in ASL that were considered as ROI. The time courses represent the average across the ten most significant vertices within each ROI in ASL. The time courses of β-power in the same vertices in the ESL (aquamarine color) are displayed for the sake of comparison. Colored horizontal bars below and above a time-course mark time intervals with the significant β-ERD and β-ERS respectively (p < 0.05, FDR-corr. for a number of time points).

To gain further insight into temporal dynamics of the learning-induced β-ERS in the ASL condition, we analyzed time courses of β-power changes in the five cortical regions of the left hemisphere, which showed a relatively early increase in the β-band power averaged across the movement execution period, i.e., before a movement was terminated. For the sake of comparison, five right-hemispheric cortical regions were also included in this analysis. The β-power time courses were derived from the most significant vertexes within each region of interest (ROI, see Methods for details). The selected cortical ROIs with the MNI coordinates of their most significant vertexes and anatomical position according to Destrieux Atlas (Destrieux et al., 2010) are given in Table 2. The location of the most significant vertexes depicted on the cortical surface in Figure 4b.

**Table 2.** Brain regions showing the β-ERS in the early post-decision period during ASL. The cortical labels derived from Destrieux Atlas (Destrieux et al., 2010). The MNI coordinates and FDR-corrected p-values for each cortical area refer to the vertex with the most significant β-ERS in the time interval 100-550 ms after movement onset.

| List of anatomical labels | The most significant vertex within each label |  |  |  |
| --- | --- | --- | --- | --- |
|  | MNI |  |  | p-value |
|  | x | y | z |  |
| Left middle-anterior part of the cingulate gyrus and sulcus | -11.13 | 18.34 | 27.38 | 0,0002 |
| Left posterior-ventral part of the cingulate gyrus (vPCC, isthmus of the cingulate gyrus) | -16.66 | -41.36 | -1.56 | 0,003 |
| Left parahippocampal gyrus (parahippocampal part of the medial occipito-temporal gyrus) | -15.53 | -41.64 | -8.84 | 0,001 |
| Left superior frontal sulcus | -17.1 | 30.66 | 43.42 | 0,0007 |
| Left inferior frontal sulcus | -39.58 | 16.65 | 22.16 | 0,00004 |
| Right middle-anterior part of the cingulate gyrus and sulcus | 10.29 | 10.09 | 43.46 | 0,006 |
| Right posterior-ventral part of the cingulate gyrus (vPCC, isthmus of the cingulate gyrus) | 17.75 | -41.29 | -4.76 | 0,002 |
| Right parahippocampal gyrus (parahippocampal part of the medial occipito-temporal gyrus) | 18.38 | -41.11 | -5.82 | 0,002 |
| Right superior frontal sulcus | 21.52 | 16.95 | 52.9 | 0,005 |
| Right inferior frontal sulcus | 42.9 | 36.66 | 4.03 | 0,007 |

For the selected ROI, the β-ERS (i.e., the significant β-power increases compared to the prestimulus baseline) was evaluated at each time point in the ASL condition. The results were principally similar to those found for the sensor-level data but revealed more information regarding the relative onset of learning-induced activity in the different cortical ROIs. The β-ERS first evolved around the motor response onset in the left prefrontal cortical areas within the anterior part of superior frontal sulcus/gyrus, the middle part of the inferior frontal sulcus/inferior frontal gyrus, dorsal part of anterior cingulate gyrus/sulcus with adjacent midline portion of SFG. Then, with a delay of more than 100 ms, the β-ERS appeared in the retrosplenial and posterior-ventral cingulate cortex, as well as in the mesial temporal structures of both hemispheres (such as parahippocampal gyrus).

### 3.4 β-ERD/β-ERS and behavioral performance – LMEM results

All participants reached the ceiling level of performance under the ASL condition, yet there was a non-random variability in response time (RT) under both ESL and ASL conditions with shorter RT signifying a better-learned response. Therefore, we chose RT as a behavioral index of the processes related to the learning of the association between auditory cues and motor responses.

For each condition and for each combined gradiometer separately, we used the linear mixedeffects model, which estimated the relationships between trial-to-trial changes in RT and β-power changes relative to the baseline while controlling for nuisance parameters such as between-subject variability and effect of body extremity used to indicate the motor response. Two sets of analyses were performed, one for the individual minimal β-power values during the Pre-movement period (β-ERD), and another one – for the individual maximal β-power values (β-ERS) during the Post-movement time interval respectively (see Methods for details).

Figure 5a demonstrates that relative changes of β-power in the Post-movement interval, but not those in the Pre-movement interval, were predicted by RT. Surprisingly though, postdecision β-power showed opposite correlations in the ESL and ASL conditions. In the exploratory learning stage (ESL), shorter RT in a given trial resulted in *greater* postmovement β-power in the gradiometers located over the posterior scalp (βmin = −0.18; all ps<0.05, FDR-corr.; see also Figure 5b). On the contrary, in the advanced learning stage, when a perfect accuracy was achieved, further shortening of reaction time led to the *smaller* post-movement β-power in gradiometers located over both posterior and anterior areas, mostly of the left hemisphere (βmax = 0.27, all ps< 0.05; FDR-corr.; see also Figure 5b).

**Figure 5.**
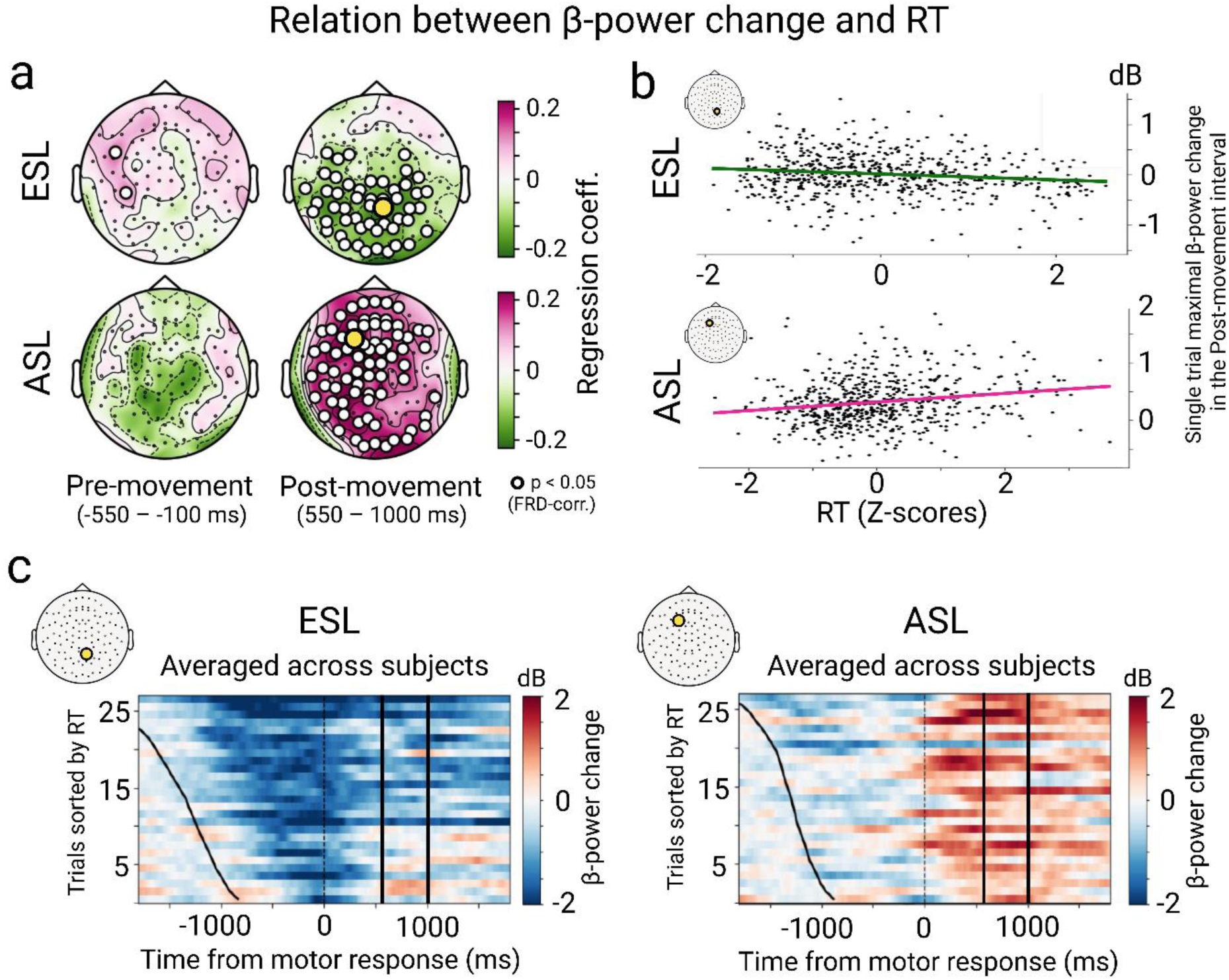
Relations between response times (RT) and β-power change in the pre- and post-movement time intervals during ESL and ASL. a) The topographic maps represent regression coefficients from the linear mixed-effects models (LMEM) performed on a single-trial basis. Greenish hue denotes negative coefficients, i.e., higher β-power values associated with shorter RT, and crimson hue – positive ones, i.e., greater β-values associated with longer RT (see Methods for details). The significant channels are marked by open white circles, and channels with the largest regression coefficient are indicated by enlarged yellow circles. b) The corresponding scatterplots from the “maximal” channels (their location is shown on the insert head) are presented for display purposes only. The x-axis represents the z-scored RT in each trial, and the y-axis -post-movement β-power change in the same trials. To control for the differences in the RT associated with movements of different limbs, the RT were z-scored for each limb separately. c) Raster plots represent the grand averages of β-power change in the “maximal” channels in single trials of each learning condition. Blue hues -β-ERD, red hues -β-ERS. For each participant, the trials were sorted in ascending order according to RT and then averaged across the participants. The vertical dotted line marks the movement onset, the left black curved lines – the stimulus onset in the respective trials, the vertical solid lines -the Post-movement time interval that was taken for LMEM in (a).

Importantly, the learning progress was accompanied by gradual shortening of RT throughout the sequential ESL and ASL learning periods (see Behavioral Results), and was paralleled by the appearance of the β-ERS towards the end of the ESL period and its further rise after the transition to ASL (Figure 5c). This suggests the greatest β-ERS values characterized the trials when a participant had already learned the rules and implemented them correctly, but the correct decision still took a relatively long time to make. Further learning accelerated decision making (reduced RT) but attenuated the post-decision β-ERS.

## 4. Discussion

We explored the dynamics of β-oscillations recorded while our participants learned, through trial and error, arbitrarily paired associations between auditory pseudowords and movements of left and right arms and legs. Our findings demonstrate the temporal and spatial organization of task-related β-oscillations undergo a major transition as the learning proceeds. Early in the learning, the deep suppression of β-oscillatory activity in multiple cortical areas occurs long before the movement onset and sustains throughout the whole behavioral trial. However, as the learning advances and participants perform the task quickly and accurately, a widespread and protracted rise in β-band power emerges around the time of motor response initiation, i.e., at the end of decision-making processes. The post-decision β-power predicts trial-by-trial response times (RT) at both stages of learning (before and after the rules become familiar), but in opposite directions. At the early learning stage, responding according to progressively more familiar associative rules is paralleled by a correlated decrease in RT and an increase in post-decision β-power. Once the rules have been learned and are implemented correctly, faster (more confident) motor responses are accompanied by a relatively attenuated rise in β-power. We will further discuss these results of the learning-induced transformation in β-oscillations in the context of the current debates regarding their functional role in memory processes.

Although numerous studies have documented the association between β-oscillations and motor behavior (Alegre et al., 2003; Pfurtscheller, Graimann, Huggins, Levine, & Schuh, 2003; Sochůrková, Rektor, Jurák, & Stančák, 2006; for review, see Cheyne, 2013), there is growing agreement regarding more general functions of β-band modulations (β-ERD and β-ERS) in cognitive tasks (Engel & Fries, 2010; Miller et al., 2018; Schmidt et al., 2019; Spitzer & Haegens, 2017). Still, in the human literature, the evidence linking β-oscillations to learning exists mainly in the domain of motor learning (Moisello et al., 2015; Nelson et al., 2017; Tan et al., 2014, 2016; Torrecillos et al., 2015). Therefore, to distinguish the learning-induced changes in β-power, that are triggered by improved movement planning and execution, from those that originate from the acquisition of the task rules, we exploited wellknown properties of effector-specific asymmetry of sensory-motor β-band modulation during movements of the left or right hand. To this end, we, first, selected the trials in which the participants correctly used their left or right hand to respond to the respective auditory pseudoword, and explored the learning effects on β-power modulations within the sensorymotor areas. Then, we examined the learning-induced changes of the large-scale β-ERD and β-ERS that were common for all four body extremities.

In accord with numerous observations in the literature (for review, see Cheyne, 2013), the effector-specific β-ERD gradually deepens during the Pre-movement period, when it is lateralized towards the hemisphere contralateral to the moving hand, becomes progressively bilateral toward movement execution, and then sustains during the dynamic phase of movement execution (Figure 2g). The β-ERD is commonly considered as an index of neural activation of the respective cortical areas involved in movement planning and execution (Doyle, Yarrow, & Brown, 2005; Kilavik, Zaepffel, Brovelli, MacKay, & Riehle, 2013). The effector-specific β-ERD is substituted by the contralaterally dominant β-ERS only after movement termination (Figure 2g), which represents the classical post-movement sensorymotor β-rebound described in many other studies (Neuper, Wörtz, & Pfurtscheller, 2006; Zaepffel, Trachel, Kilavik, & Brochier, 2013). β-rebound has been previously linked to inhibitory GABAergic activity and interpreted as an active inhibitory mechanism that “clears out” the motor program content after movement execution (for review, see Kilavik et al., 2013). Our data reveal that, although being of moderate strength, these effector-specific modulations of β-power are common for the self-paced movements of the left and right hand (Figure 2d) and for the movements at both ESL and ASL stages of auditory-motor learning when movements of each hand become uniquely associated with a certain pseudoword (Figure 2g).

In agreement with the results of recent research on motor learning (Moisello et al., 2015; Tan et al., 2016), auditory-motor associative learning does not significantly change the effector-specific β-ERD during movement preparation but augments the post-movement β-ERS (Figure 2f). This learning effect on the β-ERS cannot be accounted for by improved movement kinematics (i.e., shortening of hand movements over the course of learning, see Suppl. Figure 2), and is more likely to originate from the learning-induced changes in higherorder processes controlling movements selection rather than from fine-tuning of motor planning and execution per se. Supportive evidence for this idea comes from a study by Tan and colleagues, who explored the effector-specific β-ERS before and after exploratory adjustment in a visuomotor adaptation task (Tan et al, 2016). Their participants learned by trial-and-error how to adjust hand movements directed to the visual target, while constant perturbations in visual feedback about joystick position were introduced to trigger visuomotor adaptation. The post-movement β-ERS was significantly diminished during the first 20 trials when constant perturbations were introduced, reinstated when visuomotor adaptation occurred, but then weakened again when the visual feedback went back to normal, and the previously built model of visual-motor integration produced a high prediction error. The authors interpreted the β-ERS captured by C3/C4 EEG electrodes as a neural index of confidence in the current internal model of the visuomotor integration (but see Cao & Hu (2016) for the critique of this interpretation).

In line with these findings, one can suggest that the post-movement effector-specific β-ERS is weak at the initial stage of auditory-motor learning because the forward model is uncertain or likely to be wrong, and it is augmented when the forward model repeatedly produces the desired results so that participants are confident in its correctness. This view acknowledges that a learning-induced increase in “β-rebound” might be a downstream effect of the acquired contingency rule/forward model rather than being solely bound to the sensory-motor cortex -the only cortical region that was previously explored in the context of motor learning. The whole-brain MEG analysis allowed us to assess the broader picture of the cortical β-power changes over auditory-motor learning.

Unlike the sensory-motor β-oscillations, the β-ERD outside the sensory-motor cortex does undergo major changes over the course of learning. The large-scale β-ERD is mainly observed during the early learning stage (ESL) and is significantly reduced during the advanced stage (ASL), when association rules have been already learned by the participants (Figure 3a). It seems that the role of the β-ERD is mostly important when a subject is not aware of the rule linking auditory pseudoword to a distinct movement, and both the specific cue and cue-triggered response should be encoded and held in the working memory until the external evaluation of the outcome. In keeping with these demands, the widespread β-ERD in the ESL condition occurs after the auditory cue and then sustains over the movement initiation and execution and further up to feedback presentation (Figure 3a). The occurrence of the β-ERD in multiple cortical areas during a demanding working memory (WM) task is in accord with numerous findings, both in humans (Bočková, Chládek, Jurák, Halámek, & Rektor, 2007; Brookes et al., 2011; Hanslmayr & Staudigl, 2014) and monkeys (Kornblith et al., 2016). It is proven to reflect a neural activation that is functionally relevant to encoding and formation of memory traces (Hanslmayr, Staresina, & Bowman, 2016; Miller et al., 2018). On the other hand, weakening of the β-ERD is a usual concomitant of a decrease in working memory load (Scharinger, Soutschek, Schubert, & Gerjets, 2017) and is considered an indicator of a reduction in neural resources needed to perform the task. In support of this interpretation, the relative decrease in the strength, spread, and duration of the β-ERD in the ASL versus ESL condition is associated with shortening of the reaction time, i.e., a reduction in the decision costs (see Behavioral results).

The large-scale β-ERS is completely absent during the ESL but becomes remarkably strong during the ASL when participants have already learned the association rules and only implement them repeatedly (Figure 3a). This and other important features distinguish this learning-induced β-ERS effect from the sensory-motor β-rebound. Upon its appearance, cortical topography of the β-ERS remains the same regardless of the body side to which a moving hand belongs (Figure 3a, bottom row), and even regardless of whether the upper or lower body extremity is used to implement the motor response (Figure 3b). Besides, unlike β-rebound, it is not confined to the post-movement period, since it evolves shortly after the movement onset and coexists with the weak sensory-motor β-ERD throughout the remaining period of movement execution (Figure 3b, see also Suppl. Figure 3). Thus, rather than being driven by the movement cessation, the onset of the widespread β-ERS coincides with the end of the decision-making process and the onset of the rule implementation period. After that, the β-ERS sustains until the feedback presentation and, possibly, even further. This suggests that the emergence of the strong β-ERS in the context of learning is relevant not to rule acquisition or its retention in working memory but to some auxiliary post-decision processes, which is likely to be related to a positive appraisal of a rule-based action selection.

Recent research on β-power dynamics in the statistical learning task (Bogaerts, Richter, Landau, & Frost, 2020) also emphasizes the role of rule acquisition in the β-ERS enhancement occurring after associative learning. Their participants were asked to discover and memorize the association rule without performing any overt response. In this EEG study, learning was driven by a repeated presentation of different visual shapes organized in triplets in such a way that the order of visual shapes was kept constant within a triplet (three sequentially presented shapes) but was randomly changed between the triplets. With increased pattern repetition, the β-ERS emerged after cessation of each triplet, but was absent in response to the first visual shape in the triplet, which predicts subsequent visual stimuli. Notably, neither in that nor in our study, the learning-related β-ERS enhancement can be accounted for by the post-feedback β-power increase/reward-based processing that was previously implicated in this phenomenon (HajiHosseini & Holroyd, 2015; HajiHosseini, Hutcherson, & Holroyd, 2020; Marco-Pallarés, Münte, & Rodríguez-Fornells, 2015; Sporn, Hein, & Herrojo Ruiz, 2020; Wang, Cheung, Yee, & Tse, 2020). In our study, the β-ERS evolved long before a positive feedback presentation, while in the Bogaerts’ et al study no feedback was provided at all. Thus, in both studies, the appearance of the β-ERS over the learning task marks the success in the implementation of the newly-learnt contingency rules, when the inner appraisal of this success is based on the history of prior encounters with the same task.

The cortical topography of the β-ERS (Figure 4a) reveals involvement of the memory circuitry in the generation of learning-induced β-oscillations. The cortical sources of the evolving β-ERS are estimated in a widely distributed set of cortical areas, mainly located on the dorsal and medial surface of the prefrontal cortex including anterior cingulum, and retrosplenial-parahippocampal pathways at the medial surface of the occipital and temporal lobes (Figures 4a, Table 2). All these structures constitute the cortical nodes of the cingulatethalamo-limbic memory circle, strongly implicated in memory functions (Bubb, Kinnavane, & Aggleton, 2017). We emphasize here that this rise in β-oscillations power reflects its coordinated appearance in different nodes because of highly reliable between-node correlations in the β-ERS power from trial to trial (Figure 3e,f).

Our observation of the relatively early onset of β-ERS in the prefrontal cortex (Figure 4b) corresponds to its possible role in the rule representation (Antzoulatos & Miller, 2016; Brincat & Miller, 2015, 2016; Buschman, Denovellis, Diogo, Bullock, & Miller, 2012). Specifically, Buschman with colleagues (2012) described the rule-specific synchronization of β-oscillations between electrodes positioned in the prefrontal cortex in monkeys trained to switch between two different contingency rules connecting a cue to a motor response. Given that the power of MEG β-oscillations strongly depends on its synchronization between the local neural ensembles (da Silva, 2004), a sharp increase in the prefrontal β-ERS in our study may be related to the acquisition of contingency rules that are represented through the correlated activity of frontal neuronal populations. The linguistic nature of the rule can explain the strong left-hemispheric bias in the β-ERS (Figure 4a,b), likely reflecting a shift of the underlying cortical processes to the speech-dominant left hemisphere. Then the prefrontal β-oscillations may be transmitted to cortical and subcortical nodes of the memory circuit via the cortico-limbic-thalamic pathways reflecting reverberant network interactions that actively retain the conditional rule. This possibility is in line with multiple evidence that associates β-oscillations in cortical local field potentials (LFP) with the top-down information flow, e.g., task rules (Wang, 2010).

Crucially, the power of the post-decision β-ERS in our task correlates with behavioral reaction time, further pointing towards its functional role. When the participants’ performance reached nearly perfect accuracy, the RT measured for each trial allowed us to infer the time period that the participants used to form the correct decision. Surprisingly, the LMEM model showed that shorter RT (lower decision costs of correct response) in the ASL condition was followed by the weaker β-ERS than the β-ERS after the more lengthy/laborious correct decisions (Figure 5). Thus, despite the fact that the strength of the post-decision β-ERS progressively increases with the gradual learning of the task rules and shortening of RT, further stabilization of learning and automatization of rule implementation leads to the reduction of β-synchronization in the memory-related cortical structures. Thus, in our study, the maximal β-ERS is bound to a distinct stage of rule acquisition when the rules are already familiar but are not overlearned, therefore their implementation still causes a delay in the decision-making process. This can explain the weak β-ERS observed in the previous human and animal studies of working memory tasks – rules of which are either simple and easy to remember (for human participants) or familiar and learned over days and weeks (for rodents and monkeys). Indeed, the β-ERS during memory retention interval in WM tasks, if observed, are usually of moderate strength and length in human EEG/MEG (Deiber et al., 2007; Spitzer, Gloel, Schmidt, & Blankenburg, 2014; but see Palva et al., 2011; Proskovec, Heinrichs-Graham, & Wilson, 2019, who reported the β-ERD in the retention interval; for review, see Pavlov & Kotchoubey, 2020) or described as transient and brief β-bursts in monkey LFP (Lundqvist, Herman, & Miller, 2018; Lundqvist et al., 2016). On the other hand, opposite relationships between the post-decision β-power and RT in the ASL and ESL conditions (Figure 5) strongly suggest that the strength of β-power increase does not unequivocally index confidence in the internal appraisal of applied rules (Tan et al., 2016) or a process of “clearing out” memory traces (e.g., Schmidt et al., 2019). Since shortening of reaction time directly indicates growing confidence, the former hypothesis predicts an invariably negative correlation between the β-power and RT that actually exists only in the initial learning (Figure 5). Contrary to this hypothesis, this correlation turns positive as learning proceeds: greater β-power predicts longer RT (less confident response). The latter account is that β-bursts are a mere reflection of inhibitory processes that “clear out” the memory content after the successful usage of the acquired rule (Lundqvist et al., 2018). In this case, the emergence of the β-ERS at the late stage of learning is simply a by-product of an inhibition that suppressed neural representation of a rule when it is no longer needed. This does not explain why the post-decision β-ERS is enhanced in the trials during which implementation of the familiar rules was relatively more time-consuming.

While the investigation of β-modulation over associative learning is scarce in human studies, there is ample evidence on the learning-induced β-ERS in rodents and monkeys. The appearance of the β-ERS after successful learning was repeatedly described for a variety of associative learning tasks and for different cortical and subcortical structures of memory-related circuitry – PFC-hippocampus (Brincat & Miller, 2015), hippocampus-entorhinal cortex (Berke, Hetrick, Breck, & Greene, 2008; Igarashi, Lu, Colgin, Moser, & Moser, 2014) and cortex-basal ganglia (Leventhal et al., 2012). Moreover, Brincat and Miller (2015) showed that learning progress is accompanied by increased β-synchrony not only within the hippocampus and PFC but also between these structures, and proposed that β-oscillations are implicated in synaptic plasticity and learning of correct associations.

Most importantly, our results on the non-linear behavior of the cortical β-ERS over associative learning in humans are remarkably similar to the trial-to-trial dynamics of learning-induced β-synchronization observed in the rodents’ hippocampus during the formation of integrated contextual representation in spatial memory (Berke et al., 2008). During the first exposure to the novel environment, β-oscillations in the hippocampus are absent regardless of the time spent in the initial exploration, and appear as soon as the exploration is completed. Then, with every new encounter, place-selective synchronization in the β-range (both in local field potentials and in neuronal spiking) rapidly grows up until it reaches maximal strength and duration but then wanes, although the animal continues exploring and the hippocampal theta synchronization remains high. This and other findings led to the hypothesis that β-oscillations in the hippocampus and related cortical structures represent a reverberating mechanism that enables plasticity and supports the consolidation of newly-learned associations in the long-term memory (Brincat & Miller, 2015).

In light of the results obtained from animal studies, we speculate that the maximal level of post-decision β-power in our study is pertinent to a distinct phase of learning that requires strengthening of neural representations for newly acquired association rules in memory. However, when this phase is over, β-power falls, thus signaling that the subject can apply the rule effortlessly. Thus, the β-power is not predictive of the subject’s confidence about the choice and is only relevant in conjunction with the history of the previous encounters with the same task. This interpretation does not conflict with the “status quo” account of the β-ERS in the cognitive domain (Engel & Fries, 2010). The “status quo” hypothesis claims that the postdecision rise in the β-ERS reflects the re-activation/replay of the “good rule” representation in the working memory but in a highly protected form of synchronized network β-band activity in order to prevent its further updating. Our explanation emphasizes that the postdecision “β-state” is stabilized and amplified during learning in order to enable strengthening of the neural representations of a newly learned rule in a distributed memory network. Once the strengthening role is completed, a weak β-synchronization is sufficient to maintain a “status quo” for this association in memory

## 5. Conclusions

We have shown that over the course of auditory pseudoword-motor learning, responding according to the progressively more familiar associative rules induces the long-lasting rise of widespread cortical β-oscillations starting around the onset of cue-triggered movement. Once correct implementation of the learned rules becomes routine, faster (more confident) motor responses are accompanied by relatively attenuated β-synchronization. Our findings suggest that maximal beta activity is pertinent to a distinct stage of learning and may serve to strengthen the newly acquired association in a distributed memory network.

## Acknowledgements

We are grateful to Anna Butorina for support with data analysis, and to Ekaterina Gordeeva for valuable guidance in figure design. The research was carried out within the project of the state assignment to MSUPE. The study was implemented at the unique research facility ‘Center for Neurocognitive Research (MEG-Center)’.

## References

Alegre, M., Labarga, A., Gurtubay, I., Iriarte, J., Malanda, A., & Artieda, J. (2003). Movement-related changes in cortical oscillatory activity in ballistic, sustained and negative movements. Experimental Brain Research, 148(1), 17–25. https://doi.org/10.1007/s00221-002-1255-x

Antzoulatos, E. G., & Miller, E. K. (2016). Synchronous beta rhythms of frontoparietal networks support only behaviorally relevant representations. ELife, 5, e17822. https://doi.org/10.7554/eLife.17822

Bagiella, E., Sloan, R. P., & Heitjan, D. F. (2000). Mixed-effects models in psychophysiology. Psychophysiology, 37(1), 13–20. https://doi.org/10.1111/1469-8986.3710013

Bates, D. (2010). lme4: Linear mixed-effects models using S4 classes. R package version 0.999375-33. http://CRAN.R-Project.Org/Package=Lme4.

Berke, J. D., Hetrick, V., Breck, J., & Greene, R. W. (2008). Transient 23–30 Hz oscillations in mouse hippocampus during exploration of novel environments. Hippocampus, 18(5), 519–529. https://doi.org/10.1002/hipo.20435

Bočková, M., Chládek, J., Jurák, P., Halámek, J., & Rektor, I. (2007). Executive functions processed in the frontal and lateral temporal cortices: Intracerebral study. Clinical Neurophysiology, 118(12), 2625–2636. https://doi.org/10.1016/j.clinph.2007.07.025

Bogaerts, L., Richter, C. G., Landau, A. N., & Frost, R. (2020). Beta-Band Activity Is a Signature of Statistical Learning. The Journal of Neuroscience, 40(39), 7523 LP –7530. https://doi.org/10.1523/JNEUROSCI.0771-20.2020

Brincat, S. L., & Miller, E. K. (2015). Frequency-specific hippocampal-prefrontal interactions during associative learning. Nature Neuroscience, 18(4), 576–581. https://doi.org/10.1038/nn.3954

Brincat, S. L., & Miller, E. K. (2016). Prefrontal Cortex Networks Shift from External to Internal Modes during Learning. The Journal of Neuroscience, 36(37), 9739 LP –9754. https://doi.org/10.1523/JNEUROSCI.0274-16.2016

Brookes, M. J., Wood, J. R., Stevenson, C. M., Zumer, J. M., White, T. P., Liddle, P. F., & Morris, P. G. (2011). Changes in brain network activity during working memory tasks: A magnetoencephalography study. NeuroImage, 55(4), 1804–1815. https://doi.org/10.1016/j.neuroimage.2010.10.074

Bubb, E. J., Kinnavane, L., & Aggleton, J. P. (2017). Hippocampal–diencephalic–cingulate networks for memory and emotion: An anatomical guide. Brain and Neuroscience Advances, 1, 2398212817723443. https://doi.org/10.1177/2398212817723443

Buschman, T. J., Denovellis, E. L., Diogo, C., Bullock, D., & Miller, E. K. (2012). Synchronous Oscillatory Neural Ensembles for Rules in the Prefrontal Cortex. Neuron, 76(4), 838–846. https://doi.org/10.1016/j.neuron.2012.09.029

Cao, L., & Hu, Y.-M. (2016). Beta Rebound in Visuomotor Adaptation: Still the Status Quo? The Journal of Neuroscience, 36(24), 6365 LP –6367. https://doi.org/10.1523/JNEUROSCI.1007-16.2016

Cheyne, D. O. (2013). MEG studies of sensorimotor rhythms: A review. Experimental Neurology, 245, 27–39. https://doi.org/10.1016/j.expneurol.2012.08.030

da Silva, F. L. (2004). Functional localization of brain sources using EEG and/or MEG data: volume conductor and source models. Magnetic Resonance Imaging, 22(10), 1533–1538. https://doi.org/10.1016/j.mri.2004.10.010

Dale, A. M., Liu, A. K., Fischl, B. R., Buckner, R. L., Belliveau, J. W., Lewine, J. D., & Halgren, E. (2000). Dynamic Statistical Parametric Mapping: Combining fMRI and MEG for High-Resolution Imaging of Cortical Activity. Neuron, 26(1), 55–67. https://doi.org/10.1016/S0896-6273(00)81138-1

Deiber, M.-P., Missonnier, P., Bertrand, O., Gold, G., Fazio-Costa, L., Ibañez, V., & Giannakopoulos, P. (2007). Distinction between Perceptual and Attentional Processing in Working Memory Tasks: A Study of Phase-locked and Induced Oscillatory Brain Dynamics. Journal of Cognitive Neuroscience, 19(1), 158–172. https://doi.org/10.1162/jocn.2007.19.1.158

Destrieux, C., Fischl, B., Dale, A., & Halgren, E. (2010). Automatic parcellation of human cortical gyri and sulci using standard anatomical nomenclature. NeuroImage, 53(1), 1–15. https://doi.org/10.1016/j.neuroimage.2010.06.010

Doyle, L. M. F., Yarrow, K., & Brown, P. (2005). Lateralization of event-related beta desynchronization in the EEG during pre-cued reaction time tasks. Clinical Neurophysiology, 116(8), 1879–1888. https://doi.org/10.1016/j.clinph.2005.03.017

Engel, A. K., & Fries, P. (2010). Beta-band oscillations—signalling the status quo? Current Opinion in Neurobiology, 20(2), 156–165. https://doi.org/10.1016/j.conb.2010.02.015

Fischl, B., Salat, D. H., Busa, E., Albert, M., Dieterich, M., Haselgrove, C., … Dale, A. M. (2002). Whole Brain Segmentation: Automated Labeling of Neuroanatomical Structures in the Human Brain. Neuron, 33(3), 341–355. https://doi.org/10.1016/S0896-6273(02)00569-X

Gallistel, C. R., Fairhurst, S., & Balsam, P. (2004). The learning curve: Implications of a quantitative analysis. Proceedings of the National Academy of Sciences, 101(36), 13124–13131. https://doi.org/10.1073/pnas.0404965101

Gilbertson, T., Lalo, E., Doyle, L., Di Lazzaro, V., Cioni, B., & Brown, P. (2005). Existing Motor State Is Favored at the Expense of New Movement during 13-35 Hz Oscillatory Synchrony in the Human Corticospinal System. The Journal of Neuroscience, 25(34), 7771 LP – 7779. Retrieved from http://www.jneurosci.org/content/25/34/7771.abstract

Glasser, M. F., Coalson, T. S., Robinson, E. C., Hacker, C. D., Harwell, J., Yacoub, E., … Van Essen, D. C. (2016). A multi-modal parcellation of human cerebral cortex. Nature, 536(7615), 171–178. https://doi.org/10.1038/nature18933

Gramfort, A., Luessi, M., Larson, E., Engemann, D. A., Strohmeier, D., Brodbeck, C., … Hämäläinen, M. S. (2014). MNE software for processing MEG and EEG data. NeuroImage, 86, 446–460. https://doi.org/10.1016/j.neuroimage.2013.10.027

Gramfort, A., Luessi, M., Larson, E., Engemann, D., Strohmeier, D., Brodbeck, C., … Hämäläinen, M. (2013). MEG and EEG data analysis with MNE-Python. Frontiers in Neuroscience, 7. https://doi.org/10.3389/fnins.2013.00267

Grent-’t-Jong, T., Oostenveld, R., Jensen, O., Medendorp, W. P., & Praamstra, P. (2013). Oscillatory dynamics of response competition in human sensorimotor cortex. NeuroImage, 83, 27–34. https://doi.org/10.1016/j.neuroimage.2013.06.051

HajiHosseini, A., & Holroyd, C. B. (2015). Reward feedback stimuli elicit high-beta EEG oscillations in human dorsolateral prefrontal cortex. Scientific Reports, 5(1), 13021. https://doi.org/10.1038/srep13021

HajiHosseini, A., Hutcherson, C. A., & Holroyd, C. B. (2020). Beta oscillations following performance feedback predict subsequent recall of task-relevant information. Scientific Reports, 10(1), 15114. https://doi.org/10.1038/s41598-020-72128-x

Hanslmayr, S., Staresina, B. P., & Bowman, H. (2016). Oscillations and Episodic Memory: Addressing the Synchronization/Desynchronization Conundrum. Trends in Neurosciences, 39(1), 16–25. https://doi.org/10.1016/j.tins.2015.11.004

Hanslmayr, S., & Staudigl, T. (2014). How brain oscillations form memories — A processing based perspective on oscillatory subsequent memory effects. NeuroImage, 85, 648–655. https://doi.org/10.1016/j.neuroimage.2013.05.121

Hashimoto, H., Hasegawa, Y., Araki, T., Sugata, H., Yanagisawa, T., Yorifuji, S., & Hirata, M. (2017). Non-invasive detection of language-related prefrontal high gamma band activity with beamforming MEG. Scientific Reports, 7(1), 14262. https://doi.org/10.1038/s41598-017-14452-3

Heinrichs-Graham, E., Wilson, T. W., Santamaria, P. M., Heithoff, S. K., Torres-Russotto, D., Hutter-Saunders, J. A. L., … Gendelman, H. E. (2014). Neuromagnetic Evidence of Abnormal Movement-Related Beta Desynchronization in Parkinson’s Disease. Cerebral Cortex, 24(10), 2669–2678. https://doi.org/10.1093/cercor/bht121

Igarashi, K. M., Lu, L., Colgin, L. L., Moser, M.-B., & Moser, E. I. (2014). Coordination of entorhinal–hippocampal ensemble activity during associative learning. Nature, 510(7503), 143–147. https://doi.org/10.1038/nature13162

Jurkiewicz, M. T., Gaetz, W. C., Bostan, A. C., & Cheyne, D. (2006). Post-movement beta rebound is generated in motor cortex: Evidence from neuromagnetic recordings. NeuroImage, 32(3), 1281–1289. https://doi.org/10.1016/j.neuroimage.2006.06.005

Kelly, S. D., McDevitt, T., & Esch, M. (2009). Brief training with co-speech gesture lends a hand to word learning in a foreign language. Language and Cognitive Processes, 24(2), 313–334. https://doi.org/10.1080/01690960802365567

Kilavik, B. E., Zaepffel, M., Brovelli, A., MacKay, W. A., & Riehle, A. (2013). The ups and downs of beta oscillations in sensorimotor cortex. Experimental Neurology, 245, 15–26. https://doi.org/10.1016/j.expneurol.2012.09.014

Klepp, A., Niccolai, V., Buccino, G., Schnitzler, A., & Biermann-Ruben, K. (2015). Language–motor interference reflected in MEG beta oscillations. NeuroImage, 109, 438–448. https://doi.org/10.1016/j.neuroimage.2014.12.077

Kopell, N., Whittington, M. A., & Kramer, M. A. (2011). Neuronal assembly dynamics in the beta1 frequency range permits short-term memory. Proceedings of the National Academy of Sciences, 108(9), 3779–3784. https://doi.org/10.1073/pnas.1019676108

Kornblith, S., Buschman, T. J., & Miller, E. K. (2016). Stimulus Load and Oscillatory Activity in Higher Cortex. Cerebral Cortex, 26(9), 3772–3784. https://doi.org/10.1093/cercor/bhv182

Leventhal, D. K., Gage, G. J., Schmidt, R., Pettibone, J. R., Case, A. C., & Berke, J. D. (2012). Basal Ganglia Beta Oscillations Accompany Cue Utilization. Neuron, 73(3), 523–536. https://doi.org/10.1016/j.neuron.2011.11.032

Lundqvist, M., Herman, P., & Miller, E. K. (2018). Working Memory: Delay Activity, Yes! Persistent Activity? Maybe Not. The Journal of Neuroscience, 38(32), 7013 LP –7019. https://doi.org/10.1523/JNEUROSCI.2485-17.2018

Lundqvist, M., Rose, J., Herman, P., Brincat, S. L., Buschman, T. J., & Miller, E. K. (2016). Gamma and Beta Bursts Underlie Working Memory. Neuron, 90(1), 152–164. https://doi.org/10.1016/j.neuron.2016.02.028

Marco-Pallarés, J., Münte, T. F., & Rodríguez-Fornells, A. (2015). The role of high-frequency oscillatory activity in reward processing and learning. Neuroscience & Biobehavioral Reviews, 49, 1–7. https://doi.org/10.1016/j.neubiorev.2014.11.014

Mierau, A., Klimesch, W., & Lefebvre, J. (2017). State-dependent alpha peak frequency shifts: Experimental evidence, potential mechanisms and functional implications. Neuroscience, 360, 146–154. https://doi.org/10.1016/j.neuroscience.2017.07.037

Miller, E. K., Lundqvist, M., & Bastos, A. M. (2018). Working Memory 2.0. Neuron, 100(2), 463–475. https://doi.org/10.1016/j.neuron.2018.09.023

Moisello, C., Blanco, D., Lin, J., Panday, P., Kelly, S. P., Quartarone, A., … Ghilardi, M. F. (2015). Practice changes beta power at rest and its modulation during movement in healthy subjects but not in patients with Parkinson’s disease. Brain and Behavior, 5(10), e00374. https://doi.org/10.1002/brb3.374

Moseley, R. L., & Pulvermüller, F. (2014). Nouns, verbs, objects, actions, and abstractions: Local fMRI activity indexes semantics, not lexical categories. Brain and Language, 132, 28–42. https://doi.org/10.1016/j.bandl.2014.03.001

Nelson, A. B., Moisello, C., Lin, J., Panday, P., Ricci, S., Canessa, A., … Ghilardi, M. F. (2017). Beta Oscillatory Changes and Retention of Motor Skills during Practice in Healthy Subjects and in Patients with Parkinson’s Disease. Frontiers in Human Neuroscience, Vol. 11, p. 104. Retrieved from https://www.frontiersin.org/article/10.3389/fnhum.2017.00104

Neuper, C., Wörtz, M., & Pfurtscheller, G. (2006). ERD/ERS patterns reflecting sensorimotor activation and deactivation. In C. Neuper & W. B. T.-P. in B. R. Klimesch (Eds.), Event-Related Dynamics of Brain Oscillations (Vol. 159, pp. 211–222). https://doi.org/10.1016/S0079-6123(06)59014-4

Neuringer, A. (2002). Operant variability: Evidence, functions, and theory. Psychonomic Bulletin & Review, 9(4), 672–705. https://doi.org/10.3758/BF03196324

Niccolai, V., Klepp, A., Weissler, H., Hoogenboom, N., Schnitzler, A., & Biermann-Ruben, K. (2014). Grasping Hand Verbs: Oscillatory Beta and Alpha Correlates of Action-Word Processing. PLOS ONE, 9(9), e108059. Retrieved from https://doi.org/10.1371/journal.pone.0108059

Oldfield, R. C. (1971). The assessment and analysis of handedness: The Edinburgh inventory. Neuropsychologia, 9(1), 97–113. https://doi.org/10.1016/0028-3932(71)90067-4

Pakenham, D. O., Quinn, A. J., Fry, A., Francis, S. T., Woolrich, M. W., Brookes, M. J., & Mullinger, K. J. (2020). Post-stimulus beta responses are modulated by task duration. NeuroImage, 206, 116288. https://doi.org/10.1016/j.neuroimage.2019.116288

Palva, S., Kulashekhar, S., Hämäläinen, M., & Palva, J. M. (2011). Localization of Cortical Phase and Amplitude Dynamics during Visual Working Memory Encoding and Retention. The Journal of Neuroscience, 31(13), 5013 LP –5025. https://doi.org/10.1523/JNEUROSCI.5592-10.2011

Pavlov, Y. G., & Kotchoubey, B. (2022). Oscillatory brain activity and maintenance of verbal and visual working memory: A systematic review. Psychophysiology, 59(5), e13735. https://doi.org/10.1111/psyp.13735

Pavlova A (2022) Individual averages of beta power at ESL and ASL (left hand).figshare. https://figshare.com/articles/dataset/Individual_averages_of_beta_power_at_ESL_and_ASL_left_hand_/19692694/1. doi: 10.6084/m9.figshare.19692694

Pavlova A (2022) Individual averages of beta power at ESL and ASL (four body extremities).figshare. https://figshare.com/articles/dataset/Individual_averages_of_beta_power_at_ESL_and_ASL_four_body_extremities_/19692664/1. doi: 10.6084/m9.figshare.19692664

Pavlova A (2022) Accelerometer data for left and right hand movements.figshare. https://figshare.com/articles/dataset/Accelerometer_data_for_left_and_right_hand_movements/19692727/1. doi: 10.6084/m9.figshare.19692727

Pavlova A (2022) Individual averages of beta power during self-paced movements.figshare. https://figshare.com/articles/dataset/Individual_averages_of_beta_power_during_self-paced_movements/19692718/1. doi: 10.6084/m9.figshare.19692718

Pavlova A (2022) Individual averages of beta power at ESL and ASL (right hand).figshare. https://figshare.com/articles/dataset/Individual_averages_of_beta_power_at_ESL_and_ASL_right_hand_/19692712/1. doi: 10.6084/m9.figshare.19692712

Pavlova, A. A., Butorina, A. V., Nikolaeva, A. Y., Prokofyev, A. O., Ulanov, M. A., Bondarev, D. P., & Stroganova, T. A. (2019). Effortful verb retrieval from semantic memory drives beta suppression in mesial frontal regions involved in action initiation. Human Brain Mapping, 40(12). https://doi.org/10.1002/hbm.24624

Pfurtscheller, G., Graimann, B., Huggins, J. E., Levine, S. P., & Schuh, L. A. (2003). Spatiotemporal patterns of beta desynchronization and gamma synchronization in corticographic data during self-paced movement. Clinical Neurophysiology, 114(7), 1226–1236. https://doi.org/10.1016/S1388-2457(03)00067-1

Pfurtscheller, G., & Lopes da Silva, F.H. (1999). Event-related EEG/MEG synchronization and desynchronization: basic principles. Clinical Neurophysiology, 110(11), 1842–1857. https://doi.org/10.1016/S1388-2457(99)00141-8

Postle, N., McMahon, K. L., Ashton, R., Meredith, M., & de Zubicaray, G. I. (2008). Action word meaning representations in cytoarchitectonically defined primary and premotor cortices. NeuroImage, 43(3), 634–644. https://doi.org/10.1016/j.neuroimage.2008.08.006

Proskovec, A. L., Heinrichs-Graham, E., & Wilson, T. W. (2019). Load modulates the alpha and beta oscillatory dynamics serving verbal working memory. NeuroImage, 184, 256–265. https://doi.org/10.1016/j.neuroimage.2018.09.022

Razorenova, A. M., Chernyshev, B. V, Nikolaeva, A. Y., Butorina, A. V, Prokofyev, A. O., Tyulenev, N. B., & Stroganova, T. A. (2020). Rapid Cortical Plasticity Induced by Active Associative Learning of Novel Words in Human Adults. Frontiers in Neuroscience, Vol. 14, p. 895. Retrieved from https://www.frontiersin.org/article/10.3389/fnins.2020.00895

Scharinger, C., Soutschek, A., Schubert, T., & Gerjets, P. (2017). Comparison of the Working Memory Load in N-Back and Working Memory Span Tasks by Means of EEG Frequency Band Power and P300 Amplitude. Frontiers in Human Neuroscience, Vol. 11. Retrieved from https://www.frontiersin.org/article/10.3389/fnhum.2017.00006

Schmidt, R., Herrojo Ruiz, M., Kilavik, B. E., Lundqvist, M., Starr, P. A., & Aron, A. R. (2019). Beta Oscillations in Working Memory, Executive Control of Movement and Thought, and Sensorimotor Function. The Journal of Neuroscience, 39(42), 8231 LP –8238. https://doi.org/10.1523/JNEUROSCI.1163-19.2019

Ségonne, F., Dale, A. M., Busa, E., Glessner, M., Salat, D., Hahn, H. K., & Fischl, B. (2004). A hybrid approach to the skull stripping problem in MRI. NeuroImage, 22(3), 1060–1075. https://doi.org/10.1016/j.neuroimage.2004.03.032

Sochůrková, D., Rektor, I., Jurák, P., & Stančák, A. (2006). Intracerebral recording of cortical activity related to self-paced voluntary movements: a Bereitschaftspotential and event-related desynchronization/synchronization. SEEG study. Experimental Brain Research, 173(4), 637–649. https://doi.org/10.1007/s00221-006-0407-9

Spitzer, B., Gloel, M., Schmidt, T. T., & Blankenburg, F. (2014). Working Memory Coding of Analog Stimulus Properties in the Human Prefrontal Cortex. Cerebral Cortex, 24(8), 2229–2236. https://doi.org/10.1093/cercor/bht084

Spitzer, B., & Haegens, S. (2017). Beyond the Status Quo: A Role for Beta Oscillations in Endogenous Content (Re)Activation. ENeuro, 4(4), ENEURO.0170-17.2017. https://doi.org/10.1523/ENEURO.0170-17.2017

Sporn, S., Hein, T., & Herrojo Ruiz, M. (2020). Alterations in the amplitude and burst rate of beta oscillations impair reward-dependent motor learning in anxiety. ELife, 9, e50654. https://doi.org/10.7554/eLife.50654

Tan, H., Jenkinson, N., & Brown, P. (2014). Dynamic Neural Correlates of Motor Error Monitoring and Adaptation during Trial-to-Trial Learning. The Journal of Neuroscience, 34(16), 5678 LP –5688. https://doi.org/10.1523/JNEUROSCI.4739-13.2014

Tan, H., Wade, C., & Brown, P. (2016). Post-Movement Beta Activity in Sensorimotor Cortex Indexes Confidence in the Estimations from Internal Models. The Journal of Neuroscience, 36(5), 1516 LP –1528. https://doi.org/10.1523/JNEUROSCI.3204-15.2016

Taulu, S., Simola, J., & Kajola, M. (2005). Applications of the Signal Space Separation Method. IEEE Transactions on Signal Processing, Vol. 53, pp. 3359–3372. https://doi.org/10.1109/TSP.2005.853302

Thomson, D. J. (2000). Multitaper analysis of nonstationary and nonlinear time series data. Nonlinear and Nonstationary Signal Processing, 317–394.

Torrecillos, F., Alayrangues, J., Kilavik, B. E., & Malfait, N. (2015). Distinct Modulations in Sensorimotor Postmovement and Foreperiod β-Band Activities Related to Error Salience Processing and Sensorimotor Adaptation. The Journal of Neuroscience, 35(37), 12753 LP –12765. https://doi.org/10.1523/JNEUROSCI.1090-15.2015

Tzagarakis, C., Ince, N. F., Leuthold, A. C., & Pellizzer, G. (2010). Beta-Band Activity during Motor Planning Reflects Response Uncertainty. The Journal of Neuroscience, 30(34), 11270 LP –11277. https://doi.org/10.1523/JNEUROSCI.6026-09.2010

Wang, L., Jensen, O., van den Brink, D., Weder, N., Schoffelen, J.-M., Magyari, L., … Bastiaansen, M. (2012). Beta oscillations relate to the N400m during language comprehension. Human Brain Mapping, 33(12), 2898–2912. https://doi.org/10.1002/hbm.21410

Wang, X. (2010). Neurophysiological and Computational Principles of Cortical Rhythms in Cognition. 1195–1268. https://doi.org/10.1152/physrev.00035.2008.

Wang, Y., Cheung, H., Yee, L. T. S., & Tse, C.-Y. (2020). Feedback-related negativity (FRN) and theta oscillations: Different feedback signals for non-conform and conform decisions. Biological Psychology, 153, 107880. https://doi.org/10.1016/j.biopsycho.2020.107880

Weiss, S., & Mueller, H. (2012). “Too Many betas do not Spoil the Broth”: The Role of Beta Brain Oscillations in Language Processing. Frontiers in Psychology, Vol. 3, p. 201. Retrieved from https://www.frontiersin.org/article/10.3389/fpsyg.2012.00201

Zaepffel, M., Trachel, R., Kilavik, B. E., & Brochier, T. (2013). Modulations of EEG Beta Power during Planning and Execution of Grasping Movements. PLOS ONE, 8(3), e60060. Retrieved from https://doi.org/10.1371/journal.pone.0060060

