## Supplemental Materials for "Learning of new associations invokes a major change in modulations of cortical beta oscillations in human adults"

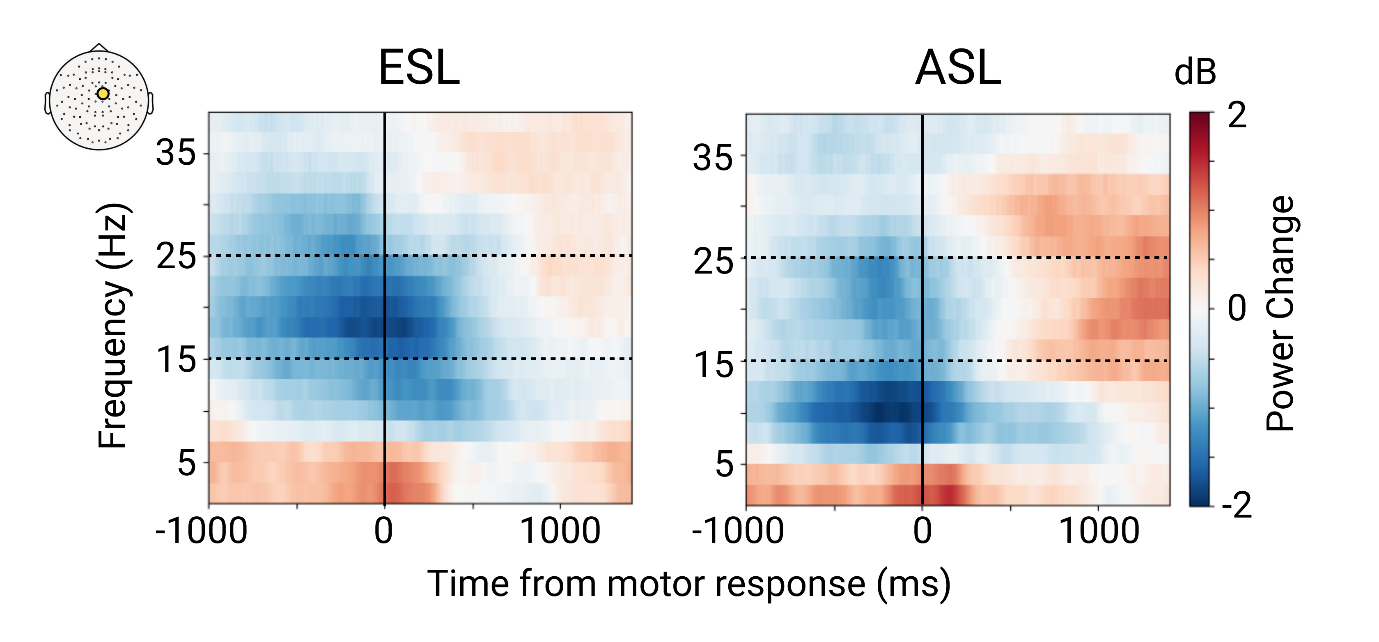

**Supplementary figure 1**. Grand average time-frequency plots from a selected sensor (marked by a yellow circle on the scalp map in the upper left corner) for motor response-related spectral power changes during the early and advanced stages of learning (ESL and ASL). All body extremities are pooled together. A vertical line marks movement onset. There is a clear learning-induced change in β-power modulation within the β-frequency range between 15 and 25 Hz (marked by dotted lines).

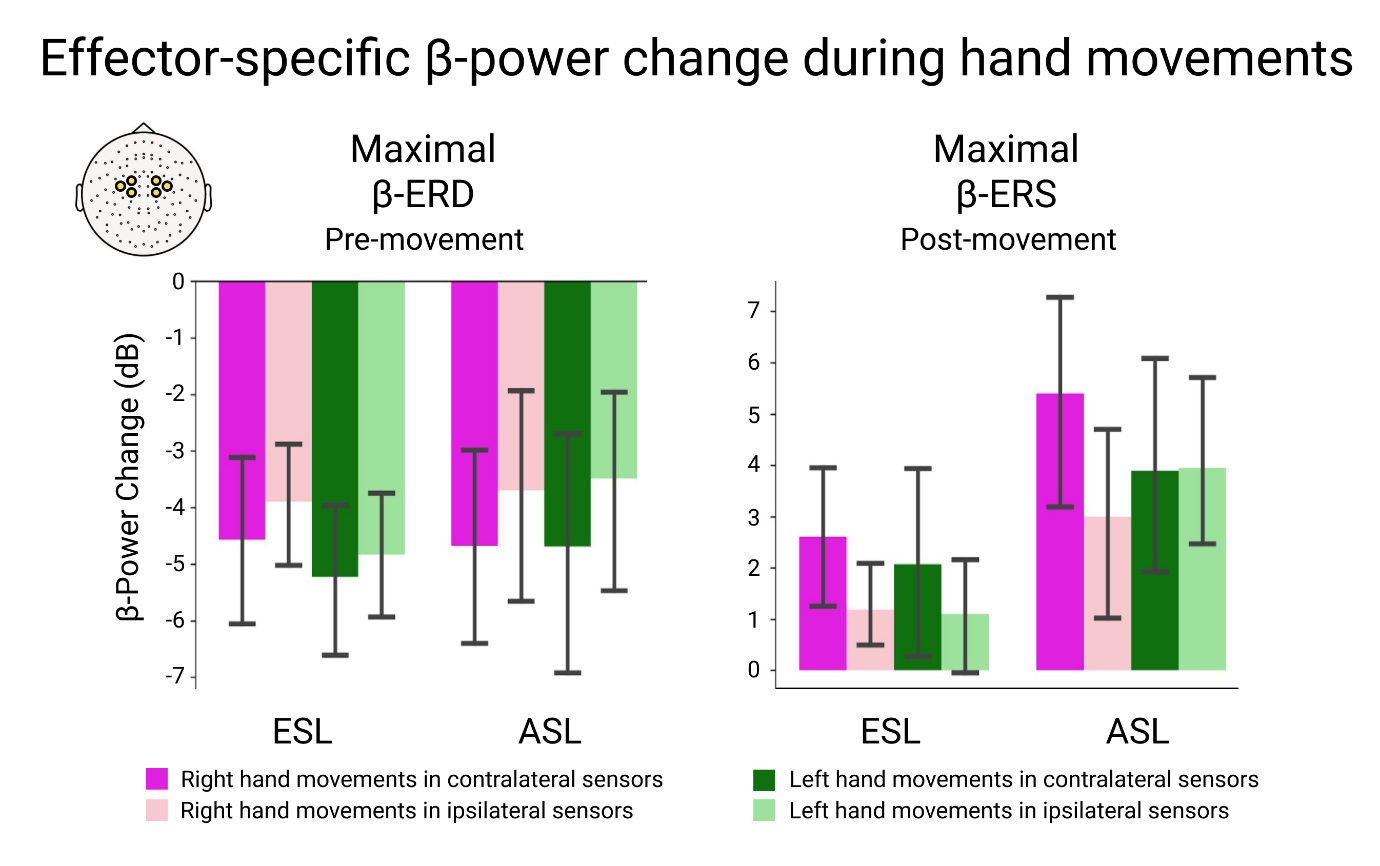

**Supplementary figure 2**. Grand average of the maximal β-ERD at Pre-movement and maximal β-ERS at Post-movement interval during contra- and ipsilateral hand movements in the subset of participants (n=13), whose accelerometer signal did not differ between the ESL and ASL conditions. Whiskers on the bar graphs indicate 95% confidence intervals.

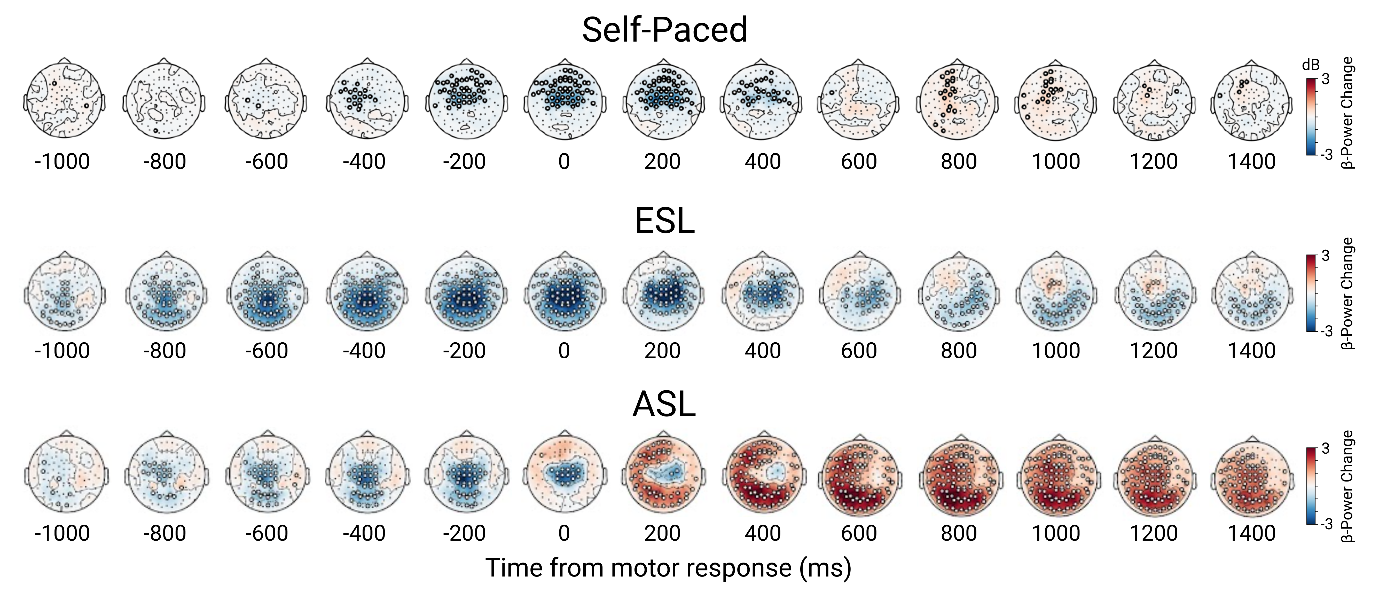

**Supplementary figure 3.** A sequence of topographic maps for β-power change (15-25 Hz) during self-paced movements and movements at the early and advanced stages of learning. Baseline normalized β-power is averaged over 50-ms intervals at 200 ms steps within -1000 to 1400 ms epochs relatively to movement onset. Movements of four extremities (left or right hand, left or right foot) were combined. Open circles indicate sensors with significant β-power changes relative to the pre-stimulus baseline (p < 0.05, FDR-corr.). Blue and red hues mark β-ERD and β-ERS respectively.

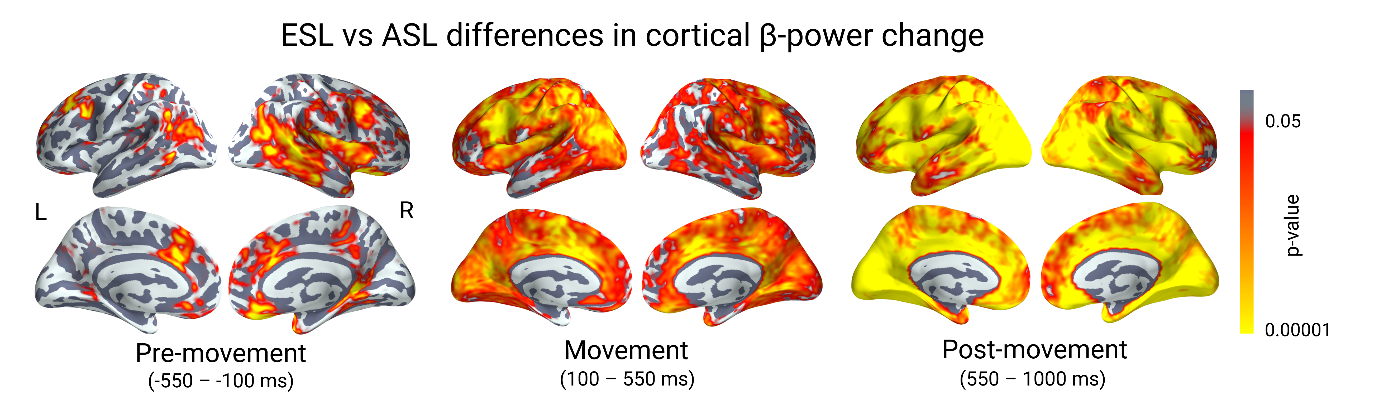

**Supplementary figure 4.** Statistical cortical maps of differences in β-power change between ESL and ASL in three successive time windows. Red-yellow hues indicate cortical regions with significant ESL versus ASL differences (p < .05, FDR-corr.).
